# Community Composition of Bacteria Isolated from Swiss Banknotes Varies Depending on Collection Environment

**DOI:** 10.1101/2021.12.01.470750

**Authors:** Anna M. Bischofberger, Alex R. Hall

## Abstract

Humans interact constantly with surfaces and associated microbial communities in the environment. The factors shaping the composition of these communities are poorly understood: some proposed explanations emphasize the influence of local habitat conditions (niche-based explanations), while others point to geographic structure and the distance among sampled locations (dispersal-based explanations). However, the relative roles of these different drivers for microbial community assembly on human-associated surfaces are not clear. Here, we used a combination of sampling, sequencing (16S rRNA) and culturing to show that the composition of banknote-associated bacterial communities varies depending on the local collection environment. Using banknotes collected from various locations and types of shops across Switzerland, we found taxonomic diversity dominated by families such as *Pseudomonadaceae* and *Staphylococcaceae*, but with banknote samples from particular types of shops (especially butcher shops) having distinct community structure. By contrast, we found no evidence of geographic structure: similarity of community composition did not decrease with increasing distance among sampled locations. These results show that microbial communities associated with banknotes, one of the most commonly encountered and exchanged human-associated surfaces, can reflect the local environmental conditions (in this case, the type of shop), and the signal for this type of variation was stronger than that for geographic structure among the locations sampled here.

## Introduction

Humans encounter microbial communities constantly and we now know that they contribute, among other things, to food and drink quality (Wareing, 2016; Reese *et al.*, 2020), soil fertility (Fierer, 2017) and human health (Libertucci and Young, 2019). Understanding variation of microbial community structure (relative abundances of different taxa) is therefore an important challenge. As for other organisms and in the wider field of community ecology, the contributions of different drivers of microbial community assembly remain unclear (Chesson, 2000; Ebach, 2015). Selection imposed by local habitat conditions and biotic interactions (niche-based explanations invoking deterministic factors) (Macarthur and Levins, 1967; Diamond, 1975) is expected to result in communities matched to their local environments. Other possible drivers include more stochastic processes, ecological drift or dispersal limitation (Hubbell, 2011), which may result in geographic structuring of communities independent from the effects of local habitat conditions. There is evidence from microbes and other taxa supporting both types of explanation (Finlay, 2002; Fargione *et al.*, 2003; Whitaker *et al.*, 2003a; Horner-Devine *et al.*, 2004; Zhou *et al.*, 2013; Webb), and it is increasingly recognised that multiple drivers are in play (Chase, 2003a; Chase and Myers, 2011; HilleRisLambers *et al.*, 2012). However, we still lack a clear picture of their relative roles (Chase, 2010; Hanson *et al.*, 2012; Stegen *et al.*, 2015; Zhou and Ning, 2017). This problem is especially acute for microbial communities on surfaces that humans interact with in daily life. For example, a recent global study of microbial communities on metro and transit systems identified city-specific community composition linked to local conditions, albeit with a shared core microbiota across locations (Danko *et al.*, 2021). However, it is not yet clear whether such biogeographical structuring is also evident on other human-associated surfaces, or how strongly such microbiomes reflect local habitat conditions (selection) as opposed to other predictors such as geographic distance among sites.

One of the most important human-associated surfaces is cash money: few everyday objects are as frequently, widely and actively exchanged among humans as banknotes. Paper-based money has been in use for hundreds of years and is now commonplace around the world. Despite the advent of credit cards and other cashless payment instruments, banknotes and coins are still the most common payment option for purchases at the physical point of sale and for person-to-person payments in the EU and Switzerland (Central Bank, 2020; Gehring *et al.*). The rough surface of most banknotes lends itself well to the attachment of microorganisms (Jalali *et al.*, 2015; Vriesekoop *et al.*, 2016), something first investigated in the late 19^th^ century (Schaarschmidt, 1884). Since then, tools such as metagenomic sequencing have enabled more detailed description of microorganisms on money, including characterization of communities on dollar bills in New York City (Maritz *et al.*, 2017), identification of pathogens and antibiotic resistance genes on Rupees in New Delhi (Jalali *et al.*, 2015), and monitoring survival of influenza viruses on banknotes (Thomas *et al.*, 2008). However, to our knowledge it remains poorly understood whether banknotes collected in different places harbour microbial communities that reflect their local environment, which would indicate a role for selection (Hanson *et al.*, 2012; Zhou and Ning, 2017), and/or show biogeographic structuring (e.g., increasing dissimilarity with geographic distance).

Here, we aimed to quantify variation of the bacterial microbiome associated with banknotes originating from different locations across a country, and to test whether this variation can be explained by geographic distances among locations, or other characteristics of the local sampling environments. We did this by collecting banknotes from each of ten locations in Switzerland (Fig. 1A), visiting five different types of shops in each location. We hypothesized community composition would vary among banknotes collected from different types of shops (if there is an effect of local habitat selection on the monetary microbiome), and/or that banknotes collected from more distant locations would harbour more dissimilar communities (if there is biogeographic structuring in general). We studied the samples isolated from the 50 banknotes by (1) 16S rRNA sequencing to determine community composition, including viable and non-viable, culturable and non-culturable bacteria, and (2) plating on chromogenic agar to detect viable, culturable bacteria. Our results show that the composition of bacterial communities isolated from banknotes – determined by both sequencing and by plating – varies depending on the local environment (shop type) in which banknotes are collected. By contrast, we find no evidence for geographic structuring of bacterial community composition across banknotes sampled in Switzerland.

**Figure 1.**
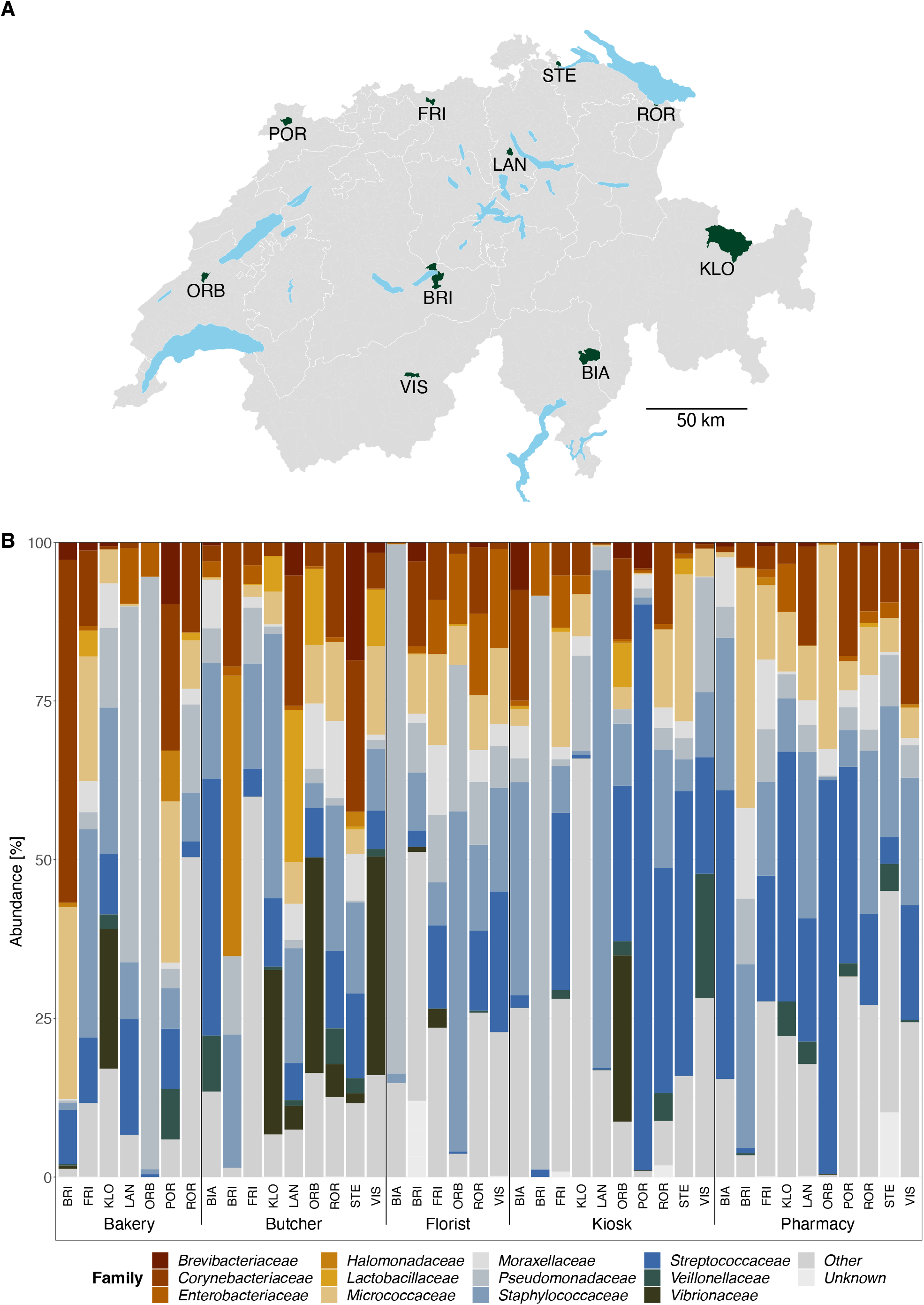
(p. 25). Bacterial community composition across sampling locations and shop types. Panel A: Distribution of sampling locations across Switzerland. The map shows the 10 municipalities, highlighted in dark green, in which banknotes were collected. Panel B: Relative abundances of the main bacterial families identified by amplicon sequencing from banknotes sampled in each shop (lower *x*-axis) and location (upper *x*-axis, indicated by three-letter abbreviations). *Unknown* contains non-assigned ZOTUs; *Other* contains assigned ZOTUs not belonging to the most abundant families. Abbreviations in both panels: BIA = Biasca, BRI = Brienz BE, FRI = Frick, KLO = Klosters-Serneus, LAN = Langnau am Albis, ORB = Orbe, POR = Porrentruy, ROR = Rorschach, STE = Stein am Rhein, VIS = Visp.

## Materials and Methods

### Sample Collection

We collected banknotes from ten locations in Switzerland. In choosing locations, we adhered to several criteria: (1) countrywide distribution of sampled locations (Fig. 1A; (2) fewer than 10’000 inhabitants per location (because we expected any signal for geographic structuring would be stronger among smaller towns due to their being, in general, less well connected with each other and with other parts of the country (Chase, 2003b)); (3) all of the targeted shop types (bakery, butcher, florist, kiosk/newsagent, and pharmacy) present at each location; (4) no significant commuter traffic between locations (according to (Federal Office of Statistics, 2020); again we expected geographic structuring of microbial communities would be stronger among less connected towns). The chosen locations were (Fig. 1A): Biasca (BIA), Brienz BE (BRI), Frick (FRI), Klosters-Serneus (KLO), Langnau am Albis (LAN), Orbe (ORB), Porrentruy (POR), Rorschach (ROR), Stein am Rhein (STE), Visp (VIS). We chose the five shop types in an attempt to represent different environments, products and clientele. More details on locations and shops can be found in Tables S1 and S2, respectivley.

On arrival in a location, we collected the banknotes from the different shops as follows: donning sterile gloves before entering the shop; choosing an item for less than CHF 10; paying the item with a CHF 20 banknote; folding the CHF 10 banknote received as change in half twice and collecting it in a sterile Nasco Whirl-Pak® sample bag (product nr. 11701973; Fisher Scientific AG, Reinach, Switzerland); storing the sample bag in a paper envelope. Once we had the five banknotes of a given location, we transported the banknotes back to the laboratory and did the initial processing as described below. All samples were collected by the same person between March 12^th^ and March 26^th^, 2021. Samples from different locations were collected on different days. One sample (butcher-BIA) was classified as “butcher” even though it came from a supermarket with a served meat counter (both butchers in Biasca (BIA) were closed on collection day); this is discussed further below. We contacted all shop owners via email to inform them about our project and gave them the option to have the banknote collected in their shop removed from the study. No shop owner vetoed inclusion of the banknote collected in their shop in the study.

### Initial Sample Processing and Plating

We performed initial processing for all banknotes on the day of collection as follows: after removing the banknote from the sample bag and placing it on a sterile surface, we swabbed the front of the banknote (not exposed to the bag surface) along the width from top to bottom (Maritz *et al.*, 2017), using a sterile cotton swab (product nr. 80.628; Sarstedt, Nürnberg, Germany) moistened with phosphate-buffered saline (PBS; product nr. P4417-100TAB; Sigma-Aldrich, Merck KgaA, Germany). Turning the banknote 90 degrees, we repeated the procedure. With the same swab, we swabbed the back of the banknote according to the same protocol. We cut the tip of the swab with sterile cissors and placed it in an 2mL Eppendorf tube filled with 950μL PBS. We repeated the procedure with the other four banknotes, sterilizing the surface between samples. As a negative control, we swabbed the interior of a sample bag and followed the protocol just described. We vortexed all Eppendorf tubes for 5min at the highest amplitude and removed the swab tips with sterile tweezers, squeezing out the remaining liquid.

To screen for viable and culturable cells, we plated a 50μL aliquot of each sample on a separate Chromatic™ MH agar plate (product nr. LF-611618; Apteq AG, Cham, Switzerland). Chromatic™ MH agar is a chromogenic agar that allows for the identification of bacteria from environmental and clinical isolates by differential colouring of colonies from different species (manufacturer’s instructions and colour guide: http://www.liofilchem.net/login/pd/ifu/11618_IFU.pdf). We incubated plates at 37°C for 24h and stored the remaining samples at −20°C until further processing as described below. Note that plating on chromogenic agar provides a relatively coarse-grained identification and only for certain taxa, but we employed it here only to test (1) for the presence of viable and culturable (i.e., colony-forming) cells and (2) whether the putative species identities of resulting colonies were consistent with taxa identified by sequence analysis. More details on colony morphology and colour can be found in Table S3.

### Genetic Analysis

We used the DNeasy® PowerLyzer® PowerSoil® Kit (product nr. 12855-100; Qiagen, Hilden, Germany) to extract bacterial DNA from thawed samples, according to the manufacturer’s protocol with slight adaptations. In brief: For each sample, we added 700μL to a PowerBead tube (positive control: 700μL PBS plus *Escherichia coli* colony from agar plate; negative control: from empty sample bag, see above) and centrifuged all PowerBead tubes at 13’000rpm for 10min at 4°C. Discarding the supernatant, we dissolved the cell pellet by adding 750μL PowerBead solution and 60μL Solution C1, and inverting the tubes 10 times. After incubating the tubes at 65°C for 10min followed by incubation at 95°C for 10min, we performed bead beating with a Bead Ruptor 24 (Omni International, Inc, Kennesaw (Georgia), United States of America (USA)) with the following settings: cycle nr = 2, strength = 5.5, time = 45s, interval = 0. After transferring the supernatant to clean 2mL collection tubes, we added 250μL Solution C2, vortexed the tubes for 5s at the highest amplitude and incubated them at 4°C for 5min. After centrifugation at 13’000rpm for 1min at 4°C, we transferred 600μL supernatant into clean 2mL collection tubes and added 200μL Solution C3. We vortexed tubes briefly at the highest amplitude and incubated them at 4°C for 5min, followed by centrifugation at 13’000 rpm for 1min at 4°C. Avoiding the pellet, we transferred 750μL supernatant into clean 2mL collection tubes, added 1200μL well-mixed Solution C4 and vortexed tubes for 5s at the highest amplitude. We loaded 675μL sample onto MB spin Columns, centrifuged them at 13’000rpm for 1min at 4°C and discarded the flow-through. We repeated this until the entire sample was processed. After adding 500μL Solution C5, we centrifuged samples at 13’000rpm for 1min at 4°C and discarded the flow-through, followed by centrifugation at 13’000rpm at 4°C for 30s. We placed the MB Spin Columns into clean 2mL collection tubes and added 70μL Solution C6 to the centre of the filter membrane. Following incubation at 65°C for 5min and centrifugation at 13’000rpm at 4°C for 1min, we discarded the MB Spin Columns and stored samples at −20°C. The 50 samples were processed in three batches (consisting of randomly selected samples) on different days, with a positive and negative control for each batch. After sequencing, we did not detect an effect of extraction batch on either read counts per sample (ANOVA: *F*_2,47_ = 2.521, *p* = 0.091) or community composition (PERMANOVA: *F*_2,41_ = 1.711, *p* > 0.1).

We quantified the obtained DNA using the Quant-iT™ dsDNA BR Assay Kit (product nr. Q33130; Thermo Fisher Scientific, Waltham, USA) in the Spark® 10M multimode microplate reader (Tecan Group Ltd, Männedorf, Switzerland). We sequenced at the Genetic Diversity Centre, Zurich, Switzerland, using the MiSeq Reagent kit v3 (600 cycle PE) (Illumina, San Diego, USA) after library preparation with the Nextera XT 96 Index kit v2, Set D (Illumina, San Diego, USA). The sequences of the four sets of primers used during library preparation (limited cycle PCR) are listed in Table S4.

### Bioinformatic Analysis

If not noted otherwise, raw sequence data were processed with USEARCH (v11.0.667_i86linux64; (Edgar, 2010). After end trimming reads (if needed) and merging them into amplicons (commands: *filter_phix*, *filter_lowc* (threshold 25), *fastx_truncate* (trim R1/R2: 25/50), *fastx_syncpairs*, *fastq_mergepairs* (min. overlap: 30; min. identity: 60%; min. merged length: 100; min. merged quality: 8)), we trimmed reads for primers (command: *search_pcr* (amplicon size range 100-600; number of mismatches: 2; coverage: full-length (no end gaps)). We used PRINSEQ-lite 0.20.4 (Schmieder and Edwards, 2011) to check quality and size filter (parameters: size range: 150-550; GC range: 30-70; min. Q mean: 20; number of Ns: 1; low complexity: dust / 30). We clustered operational taxonomic units (OTUs) at 97% using the UPARSE-OTU algorithm (number of OTUs: 2378), and used UNOISE3 algorithm to perform denoising (error-correction) of amplicon sequence variants (zero-radius OTUs, number of ZOTUs 100%: 4464), and to cluster ZOTUs at different identity levels (number of ZOTUs 99%: 3148; number of ZOTUs 98%: 2513; number of ZOTUs 97%: 2039) (Edgar, 2016b). OTU annotation was carried out using ZOTUs 100% and the SINTAX algorithm (version 11.0.667; (Edgar, 2016a)) with Silva 16S (V128) as reference database (tax filter: 0.75).

After removing positive and negative controls and mitochondrial and chloroplast sequences, and before filtering, denoised sequencing data (ZOTUs 100%) consisted of 50 samples with 14’900’848 raw reads in total, ranging from four to 6’752’341 raw reads per sample (mean number of raw reads: 266’086.57; median number of raw reads 158’243). We filtered data in two steps: (i) removing samples with low number of raw reads (<1000), resulting in the removal of eight samples from the analysis (florist-KLO, butcher-POR, florist-POR, bakery-STE, florist-STE, bakery-BIA, bakery-VIS, florist-LAN) (ii) removal of ZOTUs present at <0.1% of total sequencing depth (which can reduce the read sum for individual samples; total sequencing depth = 14’398’160). After filtering, we had 42 samples with 10’136’186 filtered reads in total, ranging from 138 filtered reads per sample for butcher-BRI to 6’167’513 filtered reads per sample for florist-BRI (mean number of filtered reads = 241’337.76; median number of filtered reads = 88’192; Fig. S1 and Table S5; numbers according to summarize_phyloseq() function in microbiome package (version 1.16.0, (Lahti and Shetty, 2012))). We had at least three samples per location and at least six samples per shop type.

The sequencing data have been deposited in the European Nucleotide Archive under the study accession number PRJEB45390 (https://www.ebi.ac.uk/ena; data will be made accessible upon publication of the study).

### Statistics

We used denoised sequences (ZOTUs 100%) for all analyses, and conducted all analyses in R (version 4.1.2).

To quantify within-sample diversity, we used the alpha() function in the microbiome package (version 1.16.0, (Lahti and Shetty, 2012)) on filtered data to calculate several different diversity indices (Shannon diversity index; Gini-Simpson index; observed richness; Chao1 index). The Shannon diversity index *H* takes into account both richness and evenness; the Gini-Simpson index gives the probabilty that two random draws from the same sample (with replacement) are not from the same type (here, the same family); the observed richness counts the number of unique species (ZOTUs 100%) present in the sample; Chao1 index is a non-parametric estimator of species richness that assumes Poisson distribution of the data (Chao, 1984; Chao and Bunge, 2002). We used analysis of variance (ANOVA) to test whether within-sample diversity varied among locations or shop types (aov() function in the stats package (R version 4.1.3; (R Core Team, 2021))).

Before analysing between-sample diversity and to account for different library sizes (Weiss *et al.*, 2017), we normalized filtered data with a regularized-logarithm (rlog() function in the DESeq2 package (version 1.34.0; (Love *et al.*, 2014))), after comparing the two available transformations offered in DESeq2 package, regularized-logarithm transformation and variance-stabilizing transformation. We chose the regularized-logarithm transformation because it (i) is recommended for smaller datasets and for cases with large ranges of sequencing depths across samples (Love *et al.*, 2014); (ii) it performed better in the meanSdPlot generated.

As a first test of whether samples from different locations and shop types differed in terms of bacterial community structure, we used permutational multivariate analysis of variance (PERMANOVA; (Anderson, 2001; Zapala and Schork, 2006)) with the adonis2() function in the vegan package (version 2.5-7; (Oksanen *et al.*, 2020)), with filtered-and-normalized data and using Bray-Curtis dissimilarity (vegdist() function) to measure differences among sampled communities. Next, to visualise any clustering among samples in terms of community composition, we performed ordination of filtered-and-normalized data with the ordinate() function in the phyloseq package (version 1.38.0; (McMurdie and Holmes, 2013)), with non-parametric multidmensional scaling (*method* = “NMDS”) and Bray-Curtis dissimilarity (*distance* = “bray”) as input parameters. We used NMDS as the ordination method because we obtained a stress value of 0.11 for our data with NMDS ordination; stress values <0.2 generally indicate good fit (Clarke, 1993). We visualised ordination results (Fig. 3) with the plot_ordination() function in the phyloseq package (version 1.38.0; (McMurdie and Holmes, 2013)). To further characterise beta diversity, we used a heatmap (plot_heatmap() function in phyloseq package version 1.38.0 (McMurdie and Holmes, 2013), relying on (Rajaram and Oono, 2010), with rlog-transformed read data, *method* = “NMDS” and *distance* = “bray”; Figure S4) and hierarchical clustering (calculated with hclust() function in the dendextend package version 1.15.2 (Galili, 2015), with a distance matrix based on rlog-transformed read data and Bray-Curtis dissimilarity index and “ward.D2” as method; Figure S5).

**Figure 2.**
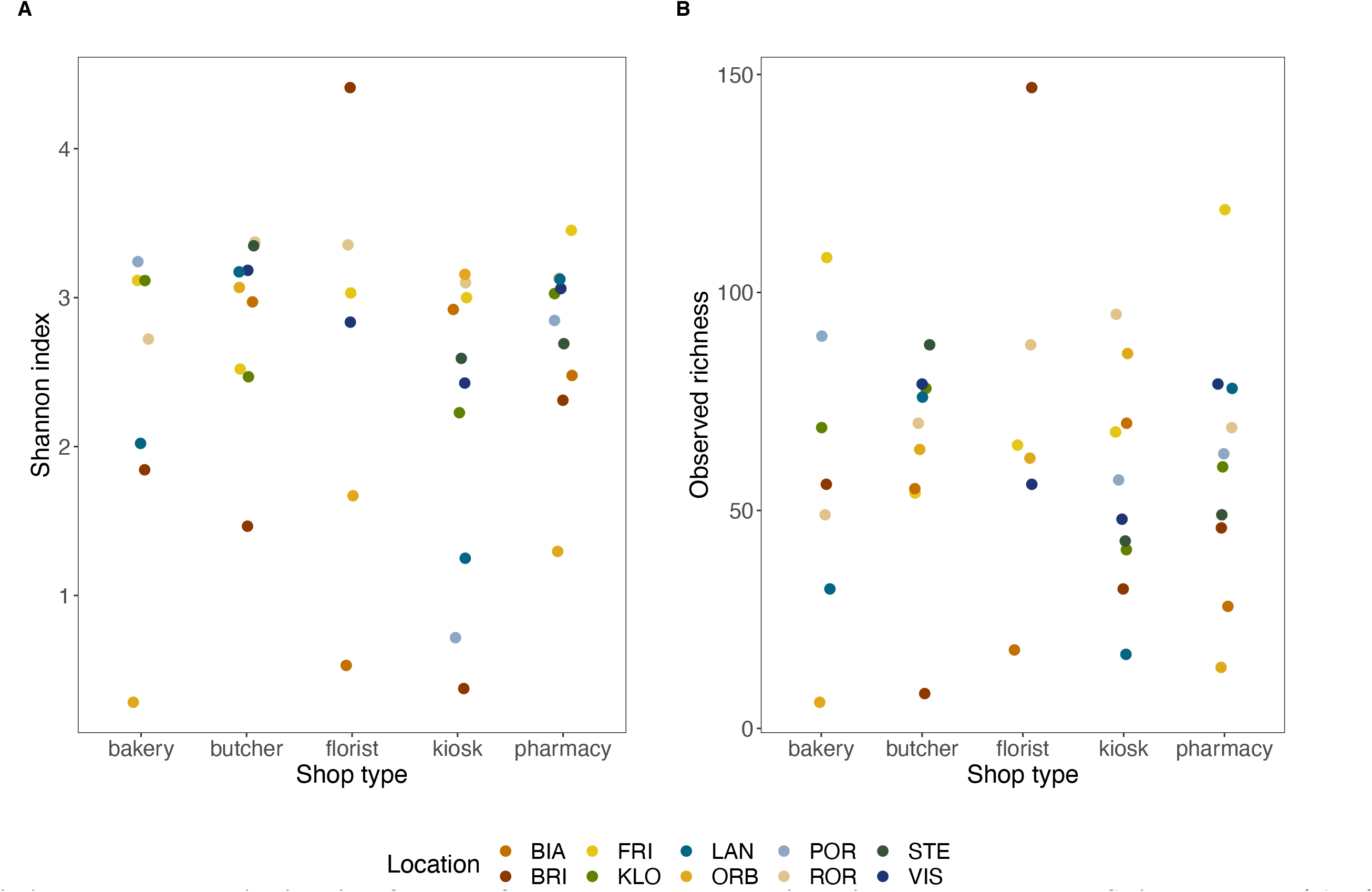
Within-sample taxonomic diversity of samples from banknotes collected in various shop types across Switzerland. Panel A: The Shannon diversity index *H* takes into account both richness (number of species present) and evenness (how evenly do present species occur). Panel B: Observed richness gives the number of unique ZOTUs present in a sample. Colours indicate locations (abbreviations: BIA = Biasca, BRI = Brienz BE, FRI = Frick, KLO = Klosters-Serneus, LAN = Langnau am Albis, ORB = Orbe, POR = Porrentruy, ROR = Rorschach, STE = Stein am Rhein, VIS = Visp). Values are jittered on the both axes in both panels.

**Figure 3.**
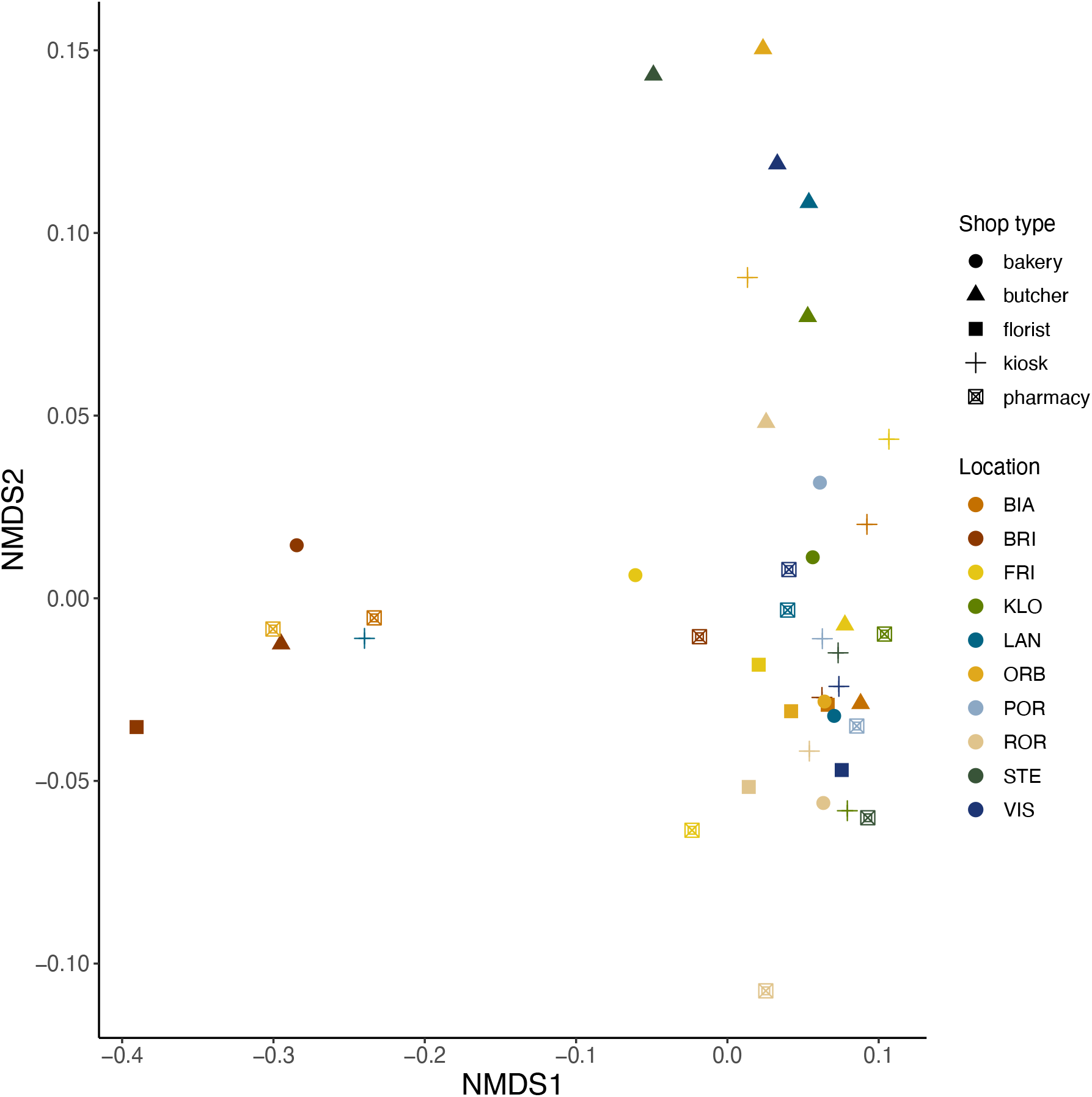
Ordination analysis. NMDS ordination based on Bray-Curtis dissimilarity between the bacterial communities isolated from banknotes (stress = 0.11). Symbols represent different shop types and are coloured according to location. Abbreviations: BIA = Biasca, BRI = Brienz BE, FRI = Frick, KLO = Klosters-Serneus, LAN = Langnau am Albis, ORB = Orbe, POR = Porrentruy, ROR = Rorschach, STE = Stein am Rhein, VIS = Visp.

We used a third type of analysis as an additional test for geographic structuring of microbial community composition, by conducting a Mantel test. This allowed us to account explicitly for the physical distance between individual sampling locations (shops), rather than testing for average differences among locations (as above in PERMANOVA). We did this using the mantel.test() function in the ape package (version 5.5; (Paradis and Schliep, 2019)), with the vegdist() function in the vegan package (version 2.5-7; (Oksanen *et al.*, 2020)) for Bray-Curtis dissimilarity of filtered-and-normalized data. We used the geodist() function in the geodist package (version 0.0.7; (Padgham and Sumner, 2021)) to calculate geographical distance (in metres) between individual sampling locations. We used the same functions when repeating the analysis for sample sets from individual shop types; here and elsewhere, we accounted for multiple testing using the Holm–Bonferroni method (sequential Bonferroni).

The core microbiome is defined as the species present above a certain threshold and shared among a set of samples (Salonen *et al.*, 2012; Shade and Handelsman, 2012). We determined the core microbiome, based on ZOTUs 100%, for each shop type as follows: Using filtered-and-normalized relative abundance data (transform() function in the microbiome package (version 1.16.0; “compositional” as input for *transform* argument) and subsetting data by shop type, we applied the core_members() function in the microbiome package (version 1.16.0; (Lahti and Shetty, 2012); *detection* = 0.001 and *prevalence* = 0.5) to filter out core ZOTUs for each shop type. Then, as a fourth analysis of how microbiome structure varies among shop types/locations, we drew a Venn diagram with the venn() function in the eulerr package (version 6.1.0) and determined ZOTU identity with the format_to_besthit() function in the microbiomeutilities package (version 1.00.16; (Lahti *et al.*, 2017)), followed by identification of core ZOTUs with reduce() and intersect() functions (base package, R version 4.1.2; (R Core Team, 2021)). We defined *shared core ZOTUs* as core ZOTUs that appeared in more than one subset (e.g. three shared core ZOTUs in the butcher and in the pharmacy subset, Fig. S2); we defined *unique core ZOTUs* as core ZOTUs that appear in only one subset (e.g. 13 unique core ZOTUs in butcher subset, Fig. S2).

To test for an association between the abundance of viable-and-culturable bacteria (presence/absence of colonies and number of colonies per plate, after plating samples on chromatic agar) and shop type and location we conducted hurdle regression for count data via maximum likelihood (hurdle() function in pscl package (version 1.5.5; (Jackman, 2020)), with formula *number.colonies ~ location* or *number.colonies ~ shop.type* and negative binomial regression (with log link) as count model). As our data contained several samples with zero viable cells we adjusted the count table with a dummy variable (dummy species present at count 1 in all samples; (Clarke *et al.*, 2006) to perform further analysis, being aware that we cannot be sure that all empty samples had zero viable cells for the same reason. We used the adjusted count table to perform PERMANOVA as described above.

The map of Switzerland with locations (Fig. 1A) was made with the mun.plot() function in RSwissMaps package (version 0.1.0.1; (Zumbach, 2019)) with the following arguments (where relevant): *bfs_id* = dt$bfs_nr, *year* = 2016, *boundaries* = “c”. Other packages and functions used in data processing and drawing figures: cowplot (version 1.1.1; (Wilke, 2020)); dplyr (version 1.0.7; (Wickham *et al.*, 2021)); ggplot2 (version 3.3.5; (Wickham, 2009)); ggpubr (version 0.4.0; (Kassambara, 2020)); microbial (version 0.0.20; (Guo and Gao, 2021)); ochre (version 1.0.0; (Allan *et al.*, 2021)); phyloseq.extended (version 0.1.1.6; (Mahendra, 2019)); phylosmith (version 1.0.6; (Smith, 2021)); ranacapa (version 0.1.0; (Kandlikar, 2021)); yarrr (version 0.1.5; (Phillips, 2017)); VennDiagram (version 1.7.0; (Chen, 2021)); vsn (version 3.62.0; (Huber *et al.*, 2002)).

## Results

### Communities Isolated from Banknotes Include *Pseudomonadaceae* and Staphylococcaceae in All Samples

After filtering the raw data in two steps (see *Methods*), we analysed 42 samples (from originally 50 samples) with variable numbers of reads per sample (Fig. S1 and Table S5). We detected some families of bacteria (Fig. 1B) in every banknote sample: *Pseudomonadaceae* and *Staphylococcaceae* were present in all samples (42/42, 100%; Table S6). These are both diverse families that include species closely associated with humans, e.g. as part of the human skin, respiratory tract or gut microbiome (Cogen *et al.*, 2008; Fierer *et al.*, 2008; Murphy and Parameswaran, 2009; Kim *et al.*, 2013; Fourquin-Gomez *et al.*, 2014; Robert-Pillot *et al.*, 2014; Rock and Donnenberg, 2014; Delanghe *et al.*, 2021; Skowron *et al.*, 2021), but have also been isolated from animal hosts and soil and aquatic environments (Cousin, 1999; Madhaiyan *et al.*, 2020). Other families were detected in significant abundance, but only in subsets of samples, such as *Vibrionaceae* and *Lactobacillaceae* (in 21 and 30 samples out of 42 respectively). If we plot the same data at the genus level, this supports the same conclusions (Fig. S3, Table S7): the most abundant genera (*Pseudomonas* and *Staphylococcus*) belonged to the top families (*Pseudomonadaceae* and *Staphylococcaceae*) and were present in all samples; *Photobacterium* (family *Vibrionaceae*) was present in 21 samples. Note despite this broad agreement, the top 12 genera (Fig. S3) did not correspond exactly to the top 12 families (Fig. 1B), because some families were represented by multiple genera. For example, the family *Micrococcaceae* is represented by three genera in Fig. S3 (*Kocuria, Micrococcus, Rothia*), and consequently the genera corresponding to *Lactobacillaceae* and *Brevibacteriaceae* (*Lactobacillus* and *Brevibacterium*) are not shown, even though they were the 13th and 14th most abundant.

The same taxa that we identified as being relatively abundant above resurfaced when we calculated the core microbiome for each shop type (taxa found in >50% of samples per shop type at abundance >0.1%). We identified seven dominant core ZOTUs (Dawson *et al.*, 2017; Cruaud *et al.*, 2020) shared between all shop types, attributed to the genera: *Corynebacterium*, *Propionibacterium*, *Pseudomonas*, *Streptococcus* (n = 2) and *Staphylococcus* (n = 2) (Fig. S2; detection threshold = 0.001, prevalence threshold = 0.5; Table S8). Thus, our sequence data revealed that a small set of ZOTUs was shared among microbial communities associated with Swiss banknotes from different shop types, but those communities were diverse and the relative abundances of individual taxa varied among samples.

Within-sample diversity estimated by Shannon diversity index (Fig. 2) did not vary significantly with either geographic location or shop type (ANOVA: effect of shop type – *F*_4,37_ = 0.844, *p* = 0.506; effect of location – *F*_9,32_ = 1.013, *p* = 0.45). We obtained the same qualitative result with an alternative diversity index (Gini-Simpson index), and the two diversity indices were highly correlated with each other (linear regression Gini-Simpson ~ Shannon: adjusted *R^2^* = 0.851). Similarly, within-sample species richness (taken as observed richness (Fig. 2B) or Chao1 index) did not vary significantly among locations or shop types (ANOVA observed richness: effect of shop type – *F*_4,37_ = 0.326, *p* = 0.859; effect of location – *F*_9,32_ = 0.801, *p* = 0.618; ANOVA Chao1: effect of shop type – *F*_4,37_ = 0.243, *p* = 0.912; effect of location – *F*_9,32_ = 1.041, *p* = 0.43) and was correlated with Shannon diversity (adjusted *R*^2^ = 0.629 for observed richness and *R^2^* = 0.478 for Chao1). Thus, none of the tested within-sample diversity measures varied significantly among shop types or geographic locations.

### Community Composition Was Specific to Shop Type, but Not Location

Bacterial community composition was more similar among banknotes collected from the same shop type than from different shop types (PERMANOVA: effect of shop type – *F*_4,41_ = 1.321, *p* = 0.03). Visualising the similarities/differences among samples by ordination analysis (non-metric multidimensional scaling NMDS) further supported variation of community composition among shop types (Fig. 3). In particular, several samples from butcher shops clustered together in the top-right corner of the ordination plot (Fig. 3). Note one sample classified as “butcher” but separate from this cluster (butcher-BIA) was collected from a supermarket with a served meat counter but separate cash desk (see *Methods*). Consistent with the variation among shop types being primarily driven by the distinctiveness of butcher shop samples, if we excluded butcher samples from the analysis we no longer found significant variation among shop types (PERMANOVA: effect of shop type, excluding butcher samples – *F*_3,32_ = 0.949, *p* = 0.584); this was not the case when re-running the analysis excluding sample sets from all other shop types, except for florist samples which resulted in marginal non-significance (PERMANOVA: effect of shop type, without florist samples – *F*_3,32_ = 1.287, *p* = 0.056). Visualising the clustering of samples using a heatmap (Fig. S4) and hierarchical clustering (Fig. S5) supported a similar pattern, with several butcher samples clustering together.

Looking at the ZOTUs making up the core microbiome for each shop type again supported butcher samples as being distinctive: the set of banknotes from butchers (n = 9) had the greatest number of core ZOTUs, both in total and unique to a single shop type (31 core ZOTUs in total, 13 unique core ZOTUs; Fig. S2). The 13 core ZOTUs unique to butchers belonged to the following genera: *Photobacterium* (n = 3; family *Vibrionaceae*), *Streptococcus* (n = 2; family *Streptococcaceae*), *Brochothrix* (n = 1; family *Listeriaceae*), *Corynebacterium* (n = 1; family *Corynebacteriaceae*), *Kocuria* (n = 1; family *Micrococcaceae*), *Lactobacillus* (n = 1; family *Lactobacillaceae*), *Lactococcus* (n = 1; family *Staphylococcaceae*), *Novosphingobium* (n = 1; family *Sphingomonadaaceae*), *Staphylococcus* (n = 1; family *Staphylococcaceae*), *Veillonella* (n = 1; family *Veillonellaceae*). This is consistent with our family-level analysis (Fig. 1), where some of the families to which these butcher-associated genera belong were over-represented among butcher samples (e.g., *Vibrionaceae*, *Lactobacillaceae*).

A comparison between core ZOTUs unique to butcher samples and NMDS scores (Fig. 3; Table S9) provides further information about the taxa that drive the among-shop-type variation: the *Veillonella* core ZOTU and one *Streptococcus* core ZOTU unique to butchers were among the ten ZOTUs with the highest positive scores for NMDS1; the *Lactococcus* and three *Photobacterium* core ZOTUs unique to butchers were among the ten ZOTUs with the highest positive scores for NMDS2 (again consistent with families *Vibrionaceae* and *Lactobacillaceae* being abundant and over-represented in butcher samples in Fig. 1). Taxa with relatively large negative scores for NMDS1 (Table S9) were mostly those with low abundance (and therefore not represented in Fig. 1B), although a *Pseudmonas* representative was among the 20 ZOTUs with the highest squared-and-rooted scores for NMDS1 (Table S9). The samples at the left of Fig. 3 are therefore poor in a combination of various taxa, but their similar position along this axis was not explained by either shop type or location (the six points include samples from all shop types and four locations).

We found no evidence that samples collected in different locations (towns) harboured different bacterial microbiomes (PERMANOVA: effect of location – *F*_9,41_ = 1.173, *p* = 0.067; Fig. 3). In a second test for geographic structuring of the sampled communities (Mantel test), we quantified the association between pairwise similarity among samples in terms of community composition and the geographic distances between sampling locations. This revealed no significant association between the two distance measures, neither in the full dataset (n = 42; Mantel test – *z* = 39’978’433, *p* = 0.751), nor when performed separately for each shop type (Mantel test, five tests total with p-values corrected for multiple testing – *p*_bakery_ = 1, *p*_butcher_ = 1; *p*_florist_ = 0.315; *p*_kiosk_ = 1; *p*_pharmacy_ = 1). There was no association between number of reads and location (ANOVA: effect of location – *F*_9,40_ = 0.836, *p* > 0.5). Thus, unlike for the variation among shop types observed above, we found no evidence that the differences in bacterial community composition among samples increased with geographic distance.

The observed variation in community composition among shop types was not explained by variable numbers of reads among samples from different shop types (Fig. S1; effect of shop type in ANOVA – *F*_4,45_ = 0.792, *p* > 0.5). Consistent with this, when we exclude the sample butcher-BRI, which had a very low read number (Fig. S1), our qualitative conclusions remain unchanged (PERMANOVA minus butcher-BRI: effect of shop type – *F*_4,40_ = 1.312, *p* = 0.034; effect of location – *F*_9,40_ = 1.054, *p* = 0.277; Mantel test comparing geographic distance and community composition, minus butcher-BRI: n = 41, *z* = 38’407’749, *p* = 0.691).

### Cultivation of Samples on Agar Supports Conclusions Drawn from Sequencing Data

When we plated aliquots of samples on Chromatic™ MH agar plates, we detected a total of 157 viable colonies across 24 of 50 plates (48%). The number of colonies per plate varied among samples (Fig. 4), and was highest on average for butcher samples (count model in hurdle regression model for shop type: butcher: *z* = 3.44, *p* < 0.001; all other shop types: *p* > 0.05; Fig. 4). There was no significant variation of the number of colonies per plate among geographic locations (count model in hurdle regression model for location: *p* > 0.05 for all locations). Several plates produced no colonies, but presence/absence of viable colonies was not predicted by either shop type or location (zero hurdle model in hurdle regression model for shop type and for location: *p* > 0.05 for all shop types and all locations).

**Figure 4.**
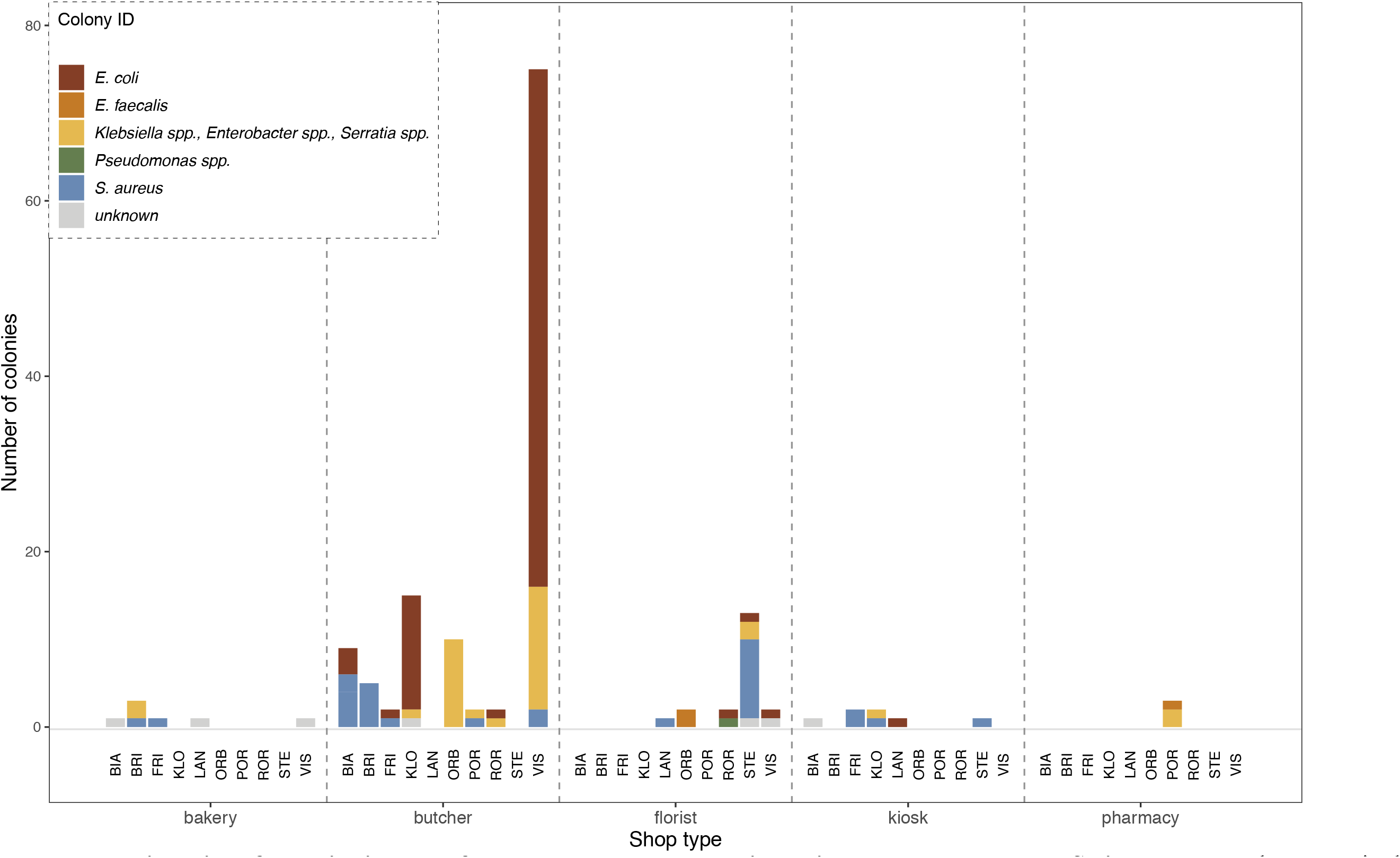
Number and identity of colonies isolated from banknotes collected in various shop types across Switzerland. Each composite bar gives the number and identity of colonies detected in samples isolated from banknotes. The bar colour indicates species identity (based on morphology and colour on Chromatic™ MH agar, see *Methods*). Colonies classified as “unknown” had uncertain identity (regular-shaped, small size) or a morphology more commonly associated with yeasts/fungi (astral appearance, polychromatic, with spores/fruiting body-like structures). Abbreviations: BIA = Biasca, BRI = Brienz BE, FRI = Frick, KLO = Klosters-Serneus, LAN = Langnau am Albis, ORB = Orbe, POR = Porrentruy, ROR = Rorschach, STE = Stein am Rhein, VIS = Visp.

The identity of colonies (as determined by colony morphology and colour on Chromatic™ MH agar plates, see *Methods*) corresponded to key taxa identified by sequencing: *E. coli = Enterobacteriaceae*; *Staphylococcus aureus = Staphlyococcaceae*; *Pseudomonas spp = Pseudomonadaceae*; *Klebsiella spp.*, *Enterobacter spp.*, *Serratia spp. = Enterobacteriaceae* (Fig. 1B Fig. 4). The exception to this trend was *Enterococcus faecalis* (3/157 colonies or 2%), which belongs to the *Enterococcaceae* family. This was not among the most abundant families in our sequence data (Fig. 1B), and we only found ZOTUs categorized as *Enterococcaceae* prior to filtering the data, indicating it was present but at lower abundances than suggested by colony formation data. For most of the other key taxa identified by sequence data that we did not find on agar plates, prior information about these taxa suggests the growth conditions (Chromatic™ MH agar; 37°C; 24h aerobic incubation) are not conducive to colony formation. For example, *Corynebacteriaceae, Lactobacillaceae*, (environmental) *Moraxellaceae*, and *Vibrionaceae* have growth optima <37°C (De Angelis and Gobbetti, 2011; Vinet and Zhedanov, 2011; Lonvaud-Funel, 2014; Tauch and Sandbote, 2014; Baron *et al.*, 2020). In addition, several of the key taxa (e.g. *Brevibacteriaceae, Corynebacteriaceae Halomonadaceae*, *Lactobacillaceae*, *Moraxellaceae*, *Vibrionaceae)* may have required incubation times >24h (Vinet and Zhedanov, 2011; Argandoña *et al.*, 2012a; Forquin and Weimer, 2014; Tauch and Sandbote, 2014; De Angelis and Gobbetti, 2016; Baron *et al.*, 2020), or different media composition such as higher salt or nutrient content (*Halomonadaceae* and *Corynebacteriaceae*, respectively, (Argandoña *et al.*, 2012b; Tauch and Sandbote, 2014)). Therefore, the absence of these taxa from our cultured samples is probably because they do not grow well in these conditions, and does not necessarily contradict our finding them in the sequence data.

As above for our community-level sequence data, the relative abundances of different taxa inferred from colony counts on chromatic agar varied among different shop types (PERMANOVA with zero-adjusted Bray-Curtis dissimilarity (Clarke *et al.*, 2006): effect of shop type – *F*_4,49_ = 2.507, *p* = 0.003) but not geographic locations (effect of location – *F*_9,41_ = 0.080, *p* = 0.701). Thus, banknotes yielded viable colonies and taxonomic identification on chromatic agar, albeit relatively coarse-grained compared to our sequence data, supporting similar conclusions (both in terms of which taxa were present and variation of community composition with collection environment but not geographic location).

## DISCUSSION

We found that Swiss banknotes taken directly from circulation harbour a variety of bacterial DNA, with the key taxa (Fig. 1B) including families found before in humans and human-associated environments, e.g. the human skin, gut or respiratory tract microbiome, or human food production (Cogen *et al.*, 2008; Fierer *et al.*, 2008; Murphy and Parameswaran, 2009; Oren, 2011; Fourquin-Gomez *et al.*, 2014; Delanghe *et al.*, 2021; Skowron *et al.*, 2021). The evidence from our data set that community composition varied depending on the shop type in which each banknote was collected is surprising: this suggests money has a microbiome reflecting its local environment, even though we assume banknotes are only transiently exposed to each habitat/shop type. By extension, we expect individual banknotes to have a dynamic microbiome that changes during their “lifecycle” as they pass from one habitat to another. By contrast, we found no evidence for geographic structuring, unlike some previous studies with microbes in other contexts (Whitaker *et al.*, 2003b; Horner-Devine *et al.*, 2004), but consistent with other observations of relatively weak biogeographic structuring in microbes (Hillebrand *et al.*, 2001; Finlay, 2002; Zhou *et al.*, 2008). Our plating results corresponded well with conclusions from sequencing, indicating that at least some taxa are present as viable, culturable cells. Together these observations suggest, on the scale investigated here, deterministic assembly processes (e.g., selection imposed by habitat heterogeneity) are more important drivers of banknote-associated bacterial community composition than other processes associated with geographic distance among sites, such as dispersal limitation (Chase, 2010).

The taxa we identified as being relatively more abundant on banknotes collected from butcher shops (*Brevibacteriaceae*, *Halomonadaceae*, *Lactobacillaceae*, *Vibrionaceae*) are known to include bacteria previously identified in “meaty” contexts. *Halomonadaceae* and *Brevibacteriaceae* have previously been found in high-salinity environments and salty foods such as poultry skin or Chinese pork cured with salt (Oren, 2011; Fourquin-Gomez *et al.*, 2014). *Brevibacteriaceae* and *Vibrionaceae* (including the genera within these families, such as *Photobacterium*, which we detected at high abundance in some butcher samples) are frequently found in samples from butchers, slaughterhouses, abattoirs (Sierra *et al.*, 1995; Pennacchia *et al.*, 2011; Połka *et al.*, 2015; Nieminen *et al.*, 2016; Fuertes-Perez *et al.*, 2019; Dourou *et al.*, 2021). Representatives of the *Lactobacillaceae* family are key species used as starter cultures in production of, and have been isolated from, cured and fermented meat (Deibel *et al.*, 1961; Samelis *et al.*, 1994; Cocolin *et al.*, 2011). This finding – that some “meaty” taxa are overrepresented in butcher samples – supports the link between collection environment and microbiome composition. However, it is important to note that members of the same families have also been isolated from a wide range of other environments including humans, plants, animals and foods (Walter, 2008; Oren, 2011; Felis and Pot, 2014; Forquin and Weimer, 2014; Franz *et al.*, 2014; Leisner and Pot, 2014; Pot *et al.*, 2014; Takemura *et al.*, 2014)(Oren, 2011). Therefore, while our results are consistent with past work finding these taxa in “meaty” environments, they can also occur elsewhere. It is interesting in this context that the sample butcher-BIA – classified as “butcher” but in fact originating from a supermarket till due to the respective location’s two butchers being closed on collection day – was relatively impoverished for the taxa linked above to the other butcher samples (butcher-BIA: 0.45% *Brevibacteriaceae*, 0.00% *Halomonadaceae*, 0.07% *Lactobacillaceae*, 0.00% *Vibrionaceae*).

At a broader level, the taxa we identified across all shop types are in line with previous research into monetary microbiomes. For example, in a metagenomic analysis of dollar bills from New York City (Maritz *et al.*, 2017) found representatives of several of the families that were dominant in our dataset (including *Staphylococcaceae*, *Micrococcaceae*, *Streptococcaceae*, *Corynebacteriaceae*, and *Moraxellaceae*). Similarly, an analysis of Chinese RMB banknotes and US dollar bills (Lin *et al.*, 2021) revealed the most abundant taxa on the family level were *Moraxellaceae*, *Pseudomonadaceae* and *Enterobacteriaceae* (all abundant in our data set). Other studies of monetary microbiomes identified similarly overlapping sets of taxa, including analysis of Indian Rupee banknotes (*Actinobacteria*, *Bacteriodetes*, *Firmicutes*, and *Proteobacteria*; no lower-taxa resolution available) (Jalali *et al.*, 2015), banknotes from Hong Kong (*Moraxellaceae*, *Micrococcaceae*, *Enterobacteriaceae, Vibrionaceae*, *Corynebacteriaceae*, *Pseudomonadaceae*, among others) (Heshiki *et al.*, 2017) and from Brazil (*Enterobacteriaceae*, *Staphylococcaceae*, *Lactobacillaceae*, *Moraxellaceae*, and *Corynebacteriaceae*, among others) (da Fonseca *et al.*, 2015). Our results go beyond past work on monetary microbiomes by showing that the relative abundances of different taxa (families and ZOTUs) vary among collection environments (shop types), but not with geographic distance within a European country.

A key implication of our results is that even temporary exposure of banknotes to certain environments is reflected by the community composition of colonizing bacteria. This is surprising because we assume the residence time of individual banknotes in a given shop is short (hours, days or weeks). It also suggests high variability in temporal beta diversity in the monetary microbiome. We found several taxa (ZOTUs) that were unique to particular shop types, and at the family level it was also visible that some taxa were overrepresented in butcher samples. Despite this, analogous to the core microbiome (Danko *et al.*, 2021) found in urban mass transit systems, we found something akin to a core microbiome across our samples in that two families (*Pseudomonadaceae, Staphylococcaceae*) were present in every sample.

Another important implication of our results is that many of the key taxa of the monetary microbiome in Switzerland are also present as viable cells (culturable on agar). This raises an interesting question about colonization dynamics in the monetary microbiome. If viable microbes are present on banknotes, are they growing and interacting on the banknote itself? We can speculate the rough surface of most banknotes not only allows attachment of bacteria, but also human skin cells, secretions and remnants of food or products during handling. Potentially, this could support bacterial growth after colonization. If the local microenvironment influences the relative growth/death rates of different taxa, this could even shape community composition and contribute to habitat-specific community composition as we observed here. Alternatively, community composition may be determined primarily at the colonization step, with different shop types harbouring different microbiomes in general (due to local habitat selection), which are “sampled” by banknotes passing through the shop. One avenue for future work aiming to tease apart these different potential drivers of the monetary microbiome would be to experimentally colonize banknotes and track community abundance and composition over time (longitudinal sampling of individual banknotes).

We emphasize strongly that our results do not address or make any predictions regarding food hygiene or potential health hazards posed to humans. We were solely interested in the microbial community composition on banknotes and how this relates to the sampling environment. Finding distinct microbiomes in different shop types does not necessarily translate to variable infection risk or indicate variable hygiene standards, only that the local monetary microbiome reflects the local commercial environment. It is, for example, not possible for us to infer how long the sampled bacteria have been present on the banknotes. One limitation of our study is that the SARS-CoV2 pandemic has led to changes in customers’ payment behaviour (Kraenzlin *et al.*, 2020). This might influence the transfer of microbiota to banknotes, because of increased hygienic awareness in customers and shop staff. For example, we observed in one case a banknote being disinfected by shop staff (the sample extracted from the banknote from this shop was removed in the initial filtering step due to low read counts, see *Methods*). However if anything we would expect any effect of pandemic-related hygiene measures to make our conclusions conservative: when hygiene standards are less stringent we might expect even more exchange of microbiota between the local environment and banknotes. In addition, further work would be needed to discover how the variation between banknotes from different shop types arises, for example, whether it is correlated with the skin microbiome of shop staff or other surfaces in the shops. Another limitation of our study is the spatial scale and number of shop types. While our study design revealed significant variation among shop types, the lack of geographic structure across Switzerland may differ from that across larger spatial scales or across different currency types.

In summary, we found that the bacterial communities associated with Swiss banknotes can vary depending on the environment in which the banknotes were collected and that, by contrast, geographic distance between sampled locations does not predict variation of the associated microbial communities. Our results also show that the most abundant taxa are not only present in the form of cell debris and genetic material, but as viable cells that can form colonies during in vitro cultivation. This raises the question of banknotes potentially being involved in transmission of bacteria between people and the environment, and/or of genes, such as antibiotic resistance genes, among bacteria co-occurring on the same banknotes. Therefore, a key avenue for future work building on the link between microbial community structure on banknotes and the local environment (shop type) identified here is to ask whether this translates to variable infection risk or horizontal gene transfer.

## Acknowledgments

AMB thanks Richard Allen, Elrina Bomben, Carole Imhof, Ricardo Leon Sampedro; Ethics Commission ETHZ, GDC ETHZ, Legal Services ETHZ, Statistical Consulting Group ETHZ. AMB and ARH thank Andrew Letten.

## Data Accessibility and Benefit-Sharing Statement

Other data presented in this study will be made available at the Dryad Digital Repository: http://dx.doi.org/...: to be completed after manuscript is accepted for publication.

Benefits Generated: Benefits from this research accrue from the sharing of our data and results on public databases as described above.

## Author Contributions

AMB, ARH designed research.

AMB performed research.

AMB, ARH analyzed data.

AMB, ARH wrote paper.

