## Supplemental Information (Tables and Figures) for "Community Composition of Bacteria Isolated from Swiss Banknotes Varies Depending on Collection Environment"

#### Table of Contents:


| | |
|---|---|
| **Supplementary Figures** | Page 2 |
| **Supplementary Tables** | Page 8 |
| **References** | Page 20 |

#### Supplementary Figures

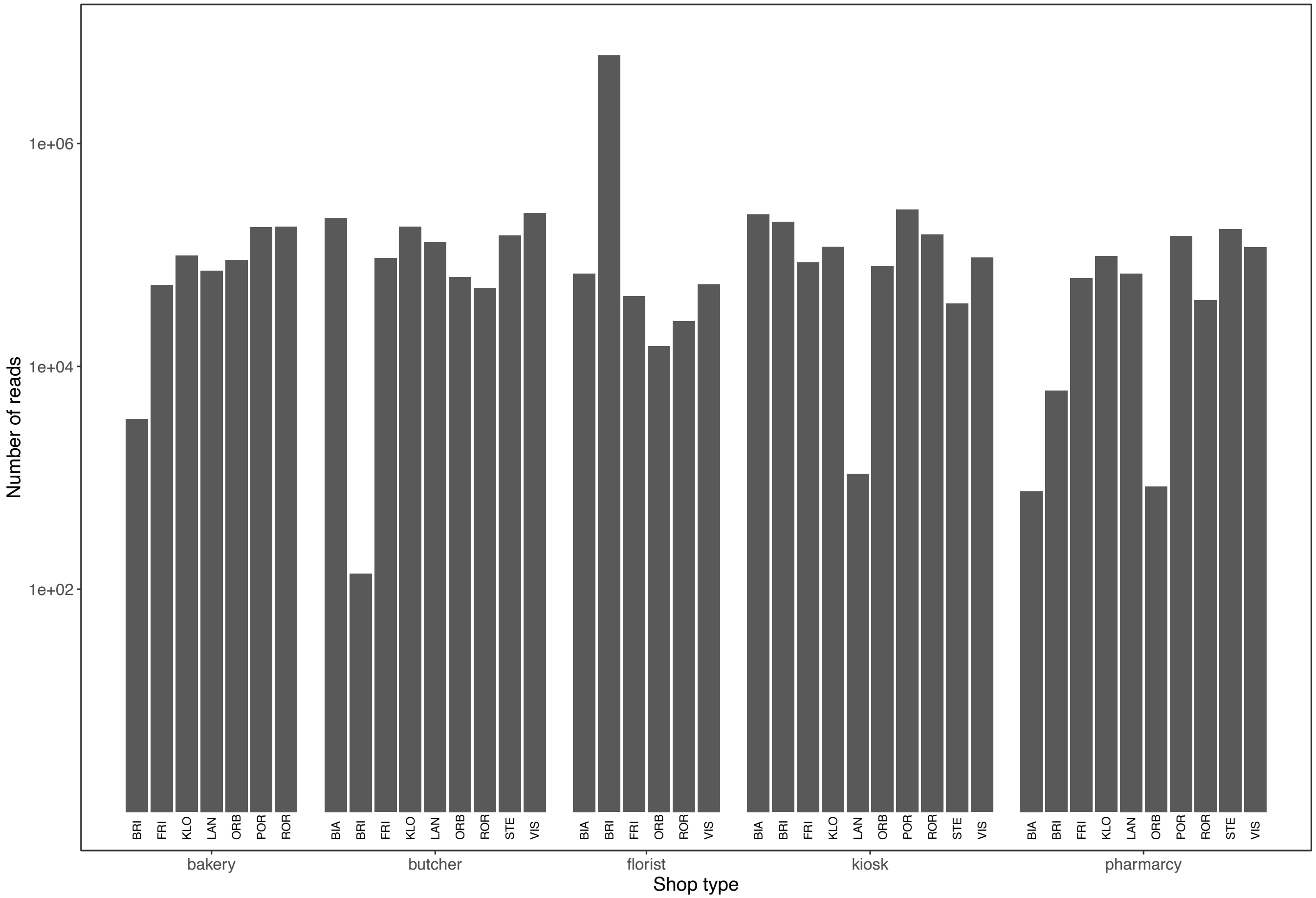


**Supplementary Figure 1. Number of reads per sample.** The figure shows the number of reads for each sample retained after filtering. Total reads per sample varied greatly, with 140 reads for the butcher in BRI and 7'152'829 reads for the florist in BRI (~51'092x difference). Note: *y*-axis is log10-transformed. Abbreviations: BIA = Biasca, BRI = Brienz BE, FRI = Frick, KLO = Klosters-Serneus, LAN = Langnau am Albis, ORB = Orbe, POR = Porrentruy, ROR = Rorschach, STE = Stein am Rhein, VIS = Visp.

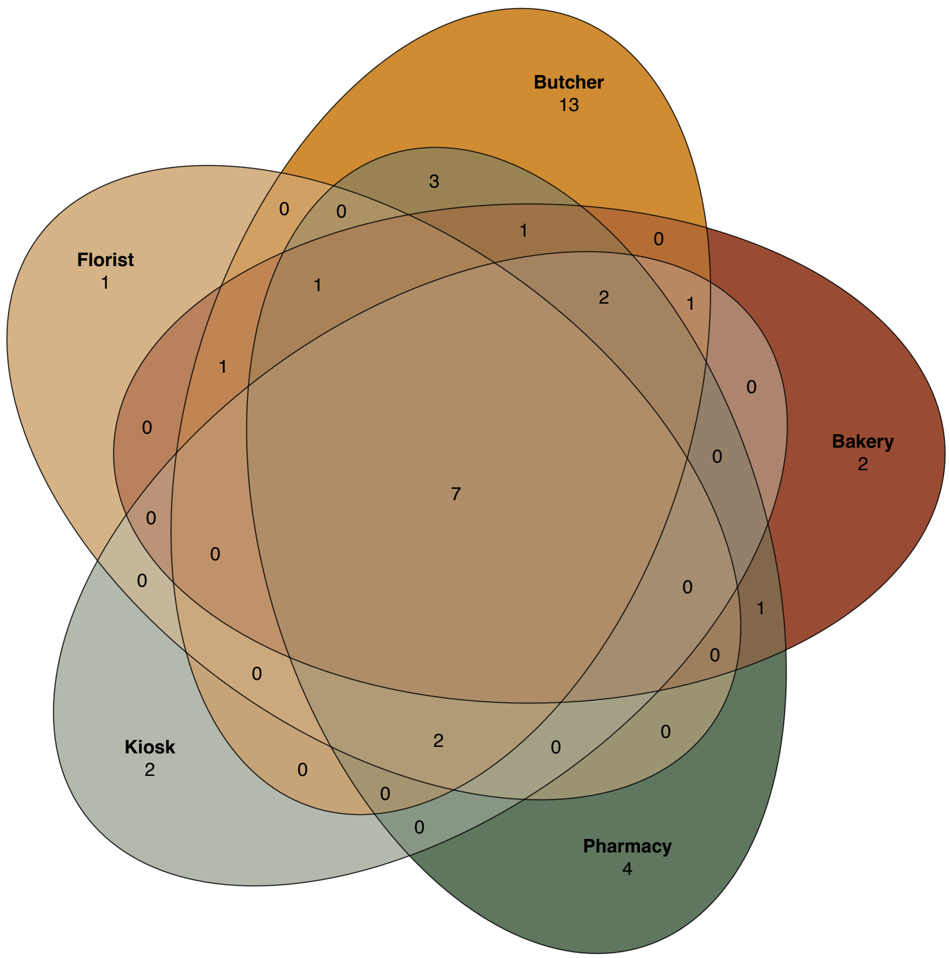


**Supplementary Figure 2. Venn diagram of unique and shared core ZOTUs in samples from banknotes collected in various shop types across Switzerland.** Each ellipse gives the number of ZOTUs present at an abundance of >0.1% in >50% of a set of banknotes attributed to a certain shop type. Numbers in overlapping areas of ellipses indicate core ZOTUs shared between sets of banknotes collected in different types of shops. The seven core microbiome ZOTUs present in >50% of every set of banknotes were attributed to the genera: *Corynebacterium*, *Escherichia-Shigella*, *Propionibacterium*, *Pseudomonas*, *Streptococcus* (n = 2), and *Staphylococcus*.

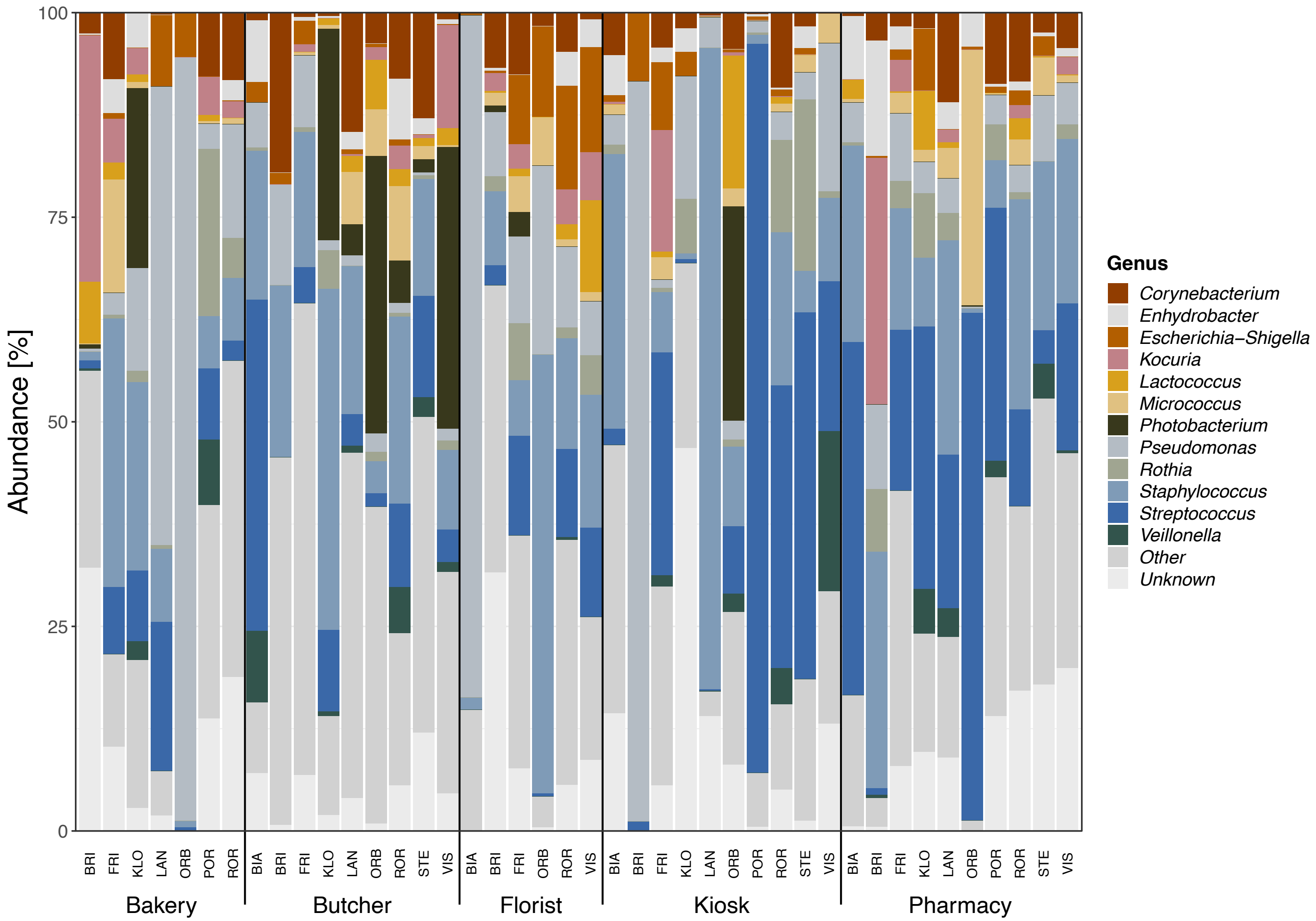


**Supplementary Figure 3. Bacterial community composition across sampling locations and shop types at Genus level.** Relative abundances of the main bacterial genera identified by amplicon sequencing from banknotes sampled in each shop (lower *x*-axis) and location (upper *x*-axis, indicated by three-letter abbreviations). *Unknown* contains non-assigned ZOTUs; *Other* contains assigned ZOTUs not belonging to the most abundant genera. The colour for genera corresponds to the colour for family in Fig. 1, except for *Kocuria* and *Rothia* which also belong to the family *Micrococcaceae.* Abbreviations: BIA = Biasca, BRI = Brienz BE, FRI = Frick, KLO = Klosters-Serneus, LAN = Langnau am Albis, ORB = Orbe, POR = Porrentruy, ROR = Rorschach, STE = Stein am Rhein, VIS = Visp.

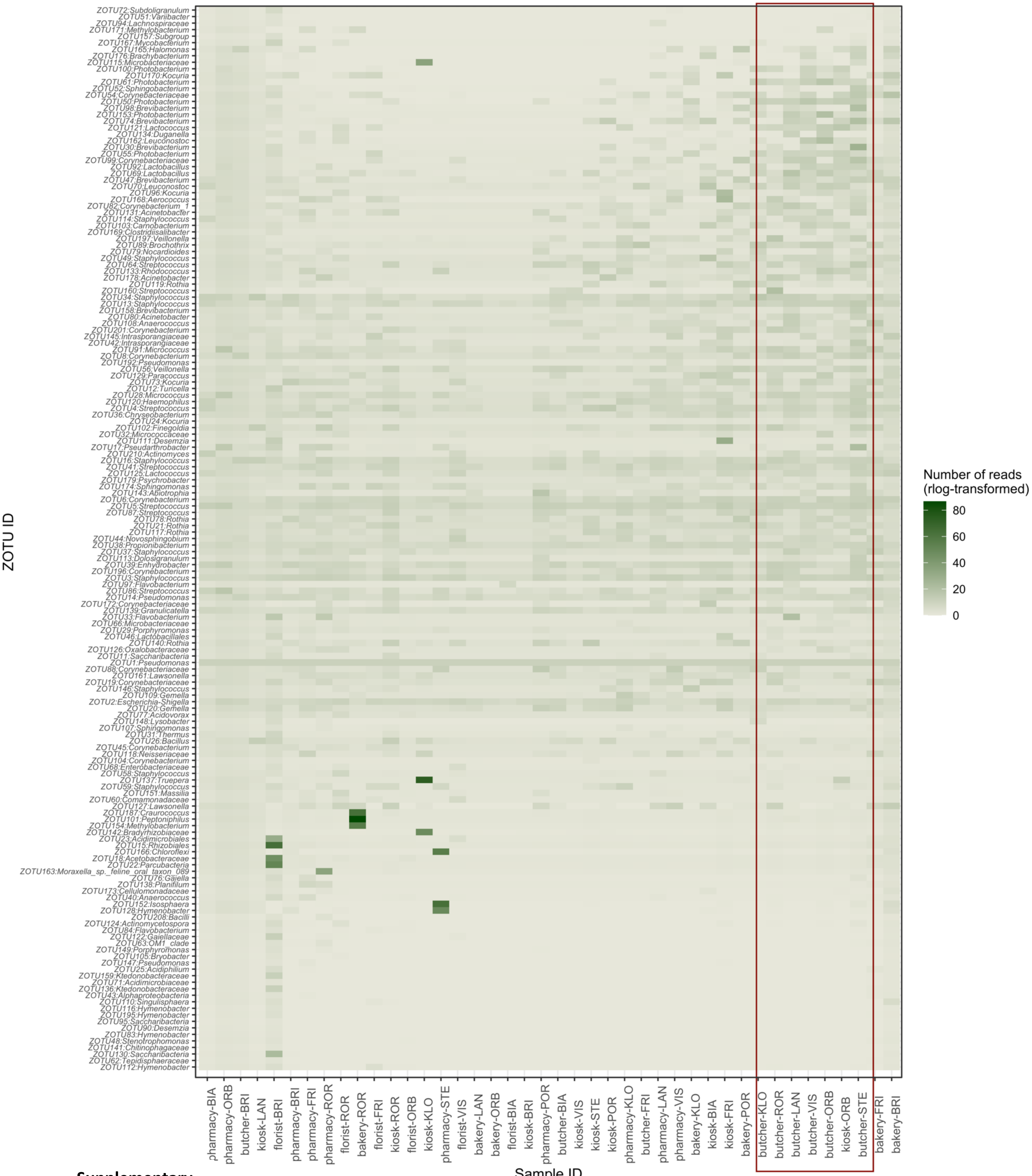


**Supplementary Figure 4.**

**Supplementary Figure 4.** Heatmap of the 163 ZOTUs occurring across samples. Darker colours indicate higher abundance of the respective ZOTU in the sample (number of reads, rlog-transformed data). The order of samples on the *x*-axis is ordination-based, i.e. more similar samples cluster together. The red rectangle comprises the six butcher samples clustering together in Fig. 3. Abbreviations: BIA = Biasca, BRI = Brienz BE, FRI = Frick, KLO = Klosters-Serneus, LAN = Langnau am Albis, ORB = Orbe, POR = Porrentruy, ROR = Rorschach, STE = Stein am Rhein, VIS = Visp.

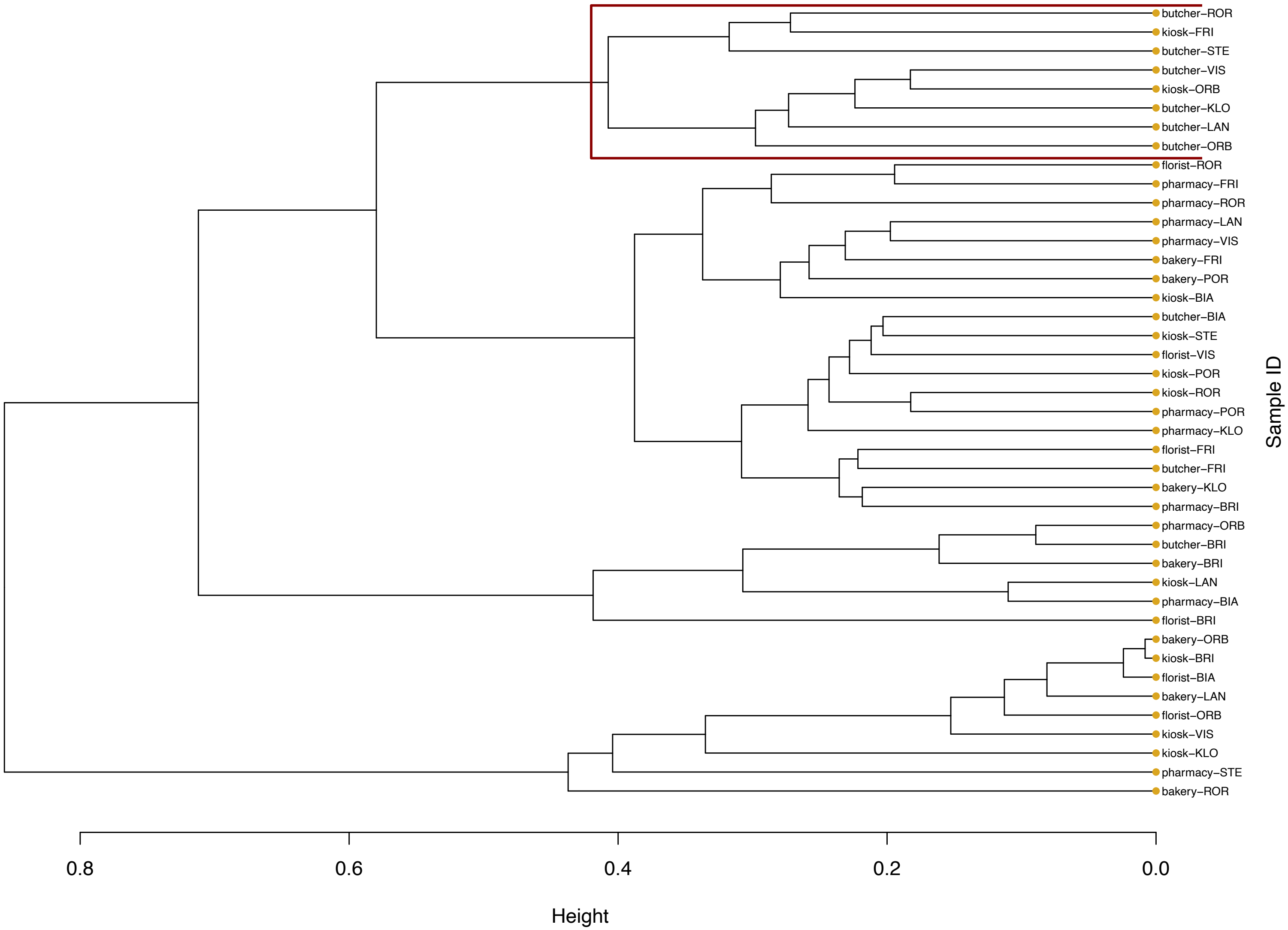

**Supplementary Figure 5.** Cluster dendogram (hierarchical clustering) of the 42 samples. The red rectangle comprises the the six butcher samples clustering toghether in Fig. 3. Abbreviations: BIA = Biasca, BRI = Brienz BE, FRI = Frick, KLO = Klosters-Serneus, LAN = Langnau am Albis, ORB = Orbe, POR = Porrentruy, ROR = Rorschach, STE = Stein am Rhein, VIS = Visp.

### Supplementary Tables

**Supplementary Table 1. Supplementary information on locations.** The tables lists supplementary information on the ten towns in Switzerland in which banknotes were collected. The linked map contains the symbols for all shops in a town from which banknotes were collected.

| Location | Location abbreviation | Canton | Postcode | Inhabitants (2018)* | Inhabitants/sqkm (2018) * | Map Notes |
|---|---|---|---|---|---|---|
| Orbe | ORB | Waadt | 1350 | 6938 | 575.8 | https://s.geo.admin.ch/8f4c69a36b |
| Porrentruy | POR | Jura | 2900 | 6678 | 452.4 | https://s.geo.admin.ch/8f9521d981 |
| Brienz | BRI | Bern | 3855 | 3092 | 64.3 | https://s.geo.admin.ch/8f4c70430c |
| Visp | VIS | Wallis | 3930 | 7950 | 602.3 | https://s.geo.admin.ch/8f4c7222fa |
| Frick | FRI | Aargau | 5070 | 5564 | 558.6 | https://s.geo.admin.ch/8f4ccde809 |
| Biasca | BIA | Tessin | 6710 | 6115 | 103.5 | https://s.geo.admin.ch/8f620d5170 |
| Klosters | KLO | Graubünden | 7250 | 4451 | 20.3 | https://s.geo.admin.ch/8f4c7a3ba8 |
| Langnau a. Albis | LAN | Zürich | 8135 | 7547 | 870.5 | https://s.geo.admin.ch/8f4c7be11d |
| Stein a. Rhein | STE | Schaffhausen | 8260 | 3415 | 590.8 | https://s.geo.admin.ch/8f8ac5922b |
| Rorschach | ROR | St Gallen | 9400 | 9441 | 5303.9 | https://s.geo.admin.ch/8f4c7f934b |
| * source: https://www.bfs.admin.ch/bfs/de/home/statistiken/regionalstatistik/regionale-portraets-kennzahlen/gemeinden/gemeindeportraets.html | | | | | | |

**Supplementary Table 2. Supplementary information on samples.** The tables lists supplementary information on the fifty banknote samples collected for this study. The column headings are: Denomination = banknotes value ; Series = number of Swiss banknotes series ; Issue year = year banknote was issued ; Collection date/hour = date/hour of banknote collection ; Extraction day/batch = day/batch of DNA extraction ; Location = town banknote was collected in ; Lat/Long_WGS84 = coordinates in WGS format; Shop type = shop type banknote was collected in. The location abbreviations are: BIA = Biasca, BRI = Brienz BE, FRI = Frick, KLO = Klosters-Serneus, LAN = Langnau am Albis, ORB = Orbe, POR = Porrentruy, ROR = Rorschach, STE = Stein am Rhein, VIS = Visp.

| Sample ID | Denomi-nation | Series | Issue year | Collection date | Collection hour | Extraction day | Extraction batch | Location | Lat_WGS84 | Long_WGS84 | Shop type |
|---|---|---|---|---|---|---|---|---|---|---|---|
| bakery-ORB | 10 | 9 | 2016 | 03.17.2021 | 10:00:00 | 03.30.2021 | 2 | ORB | 6.53214199 | 46.725701 | bakery |
| butcher-ORB | 10 | 9 | 2016 | 03.17.2021 | 10:00:00 | 03.29.2021 | 1 | ORB | 6.5319776 | 46.7245842 | butcher |
| florist-ORB | 10 | 9 | 2016 | 03.17.2021 | 10:00:00 | 03.30.2021 | 2 | ORB | 6.53205872 | 46.7236492 | florist |
| kiosk-ORB | 10 | 9 | 2016 | 03.17.2021 | 10:00:00 | 03.29.2021 | 1 | ORB | 6.53893532 | 46.7215536 | kiosk |
| pharmacy-ORB | 10 | 9 | 2016 | 03.17.2021 | 10:00:00 | 04.01.21 | 3 | ORB | 6.53241747 | 46.7233192 | pharmacy |
| bakery-POR | 10 | 9 | 2016 | 03.25.2021 | 10:00:00 | 04.01.21 | 3 | POR | 7.07815716 | 47.4210245 | bakery |
| butcher-POR | 10 | 9 | 2016 | 03.25.2021 | 10:00:00 | 03.29.2021 | 1 | POR | 7.07322855 | 47.4189223 | butcher |
| florist-POR | 10 | 9 | 2016 | 03.25.2021 | 10:00:00 | 03.30.2021 | 2 | POR | 7.0747097 | 47.417425 | florist |
| kiosk-POR | 10 | 9 | 2016 | 03.25.2021 | 11:00:00 | 03.30.2021 | 2 | POR | 7.07743321 | 47.4163902 | kiosk |
| pharmacy-POR | 10 | 9 | 2016 | 03.25.2021 | 10:00:00 | 03.29.2021 | 1 | POR | 7.07487368 | 47.416697 | pharmacy |
| bakery-BRI | 10 | 9 | 2016 | 03.22.2021 | 10:00:00 | 03.30.2021 | 2 | BRI | 8.03353591 | 46.754099 | bakery |
| butcher-BRI | 10 | 9 | 2017 | 03.22.2021 | 10:00:00 | 03.30.2021 | 2 | BRI | 8.03155907 | 46.7540553 | butcher |
| florist-BRI | 10 | 9 | 2017 | 03.22.2021 | 10:00:00 | 04.01.21 | 3 | BRI | 8.03082675 | 46.754113 | florist |
| kiosk-BRI | 10 | 9 | 2017 | 03.22.2021 | 10:00:00 | 03.29.2021 | 1 | BRI | 8.03893735 | 46.7549073 | kiosk |
| pharmacy-BRI | 10 | 9 | 2016 | 03.22.2021 | 10:00:00 | 03.29.2021 | 1 | BRI | 8.03229357 | 46.7541954 | pharmacy |
| bakery-VIS | 10 | 9 | 2016 | 03.16.2021 | 10:00:00 | 03.29.2021 | 1 | VIS | 7.88191502 | 46.2931997 | bakery |
| butcher-VIS | 10 | 9 | 2016 | 03.16.2021 | 10:00:00 | 04.01.21 | 3 | VIS | 7.88126083 | 46.2925456 | butcher |
| florist-VIS | 10 | 9 | 2016 | 03.16.2021 | 10:00:00 | 03.30.2021 | 2 | VIS | 7.88144706 | 46.2915104 | florist |
| kiosk-VIS | 10 | 9 | 2017 | 03.16.2021 | 10:00:00 | 03.30.2021 | 2 | VIS | 7.88133545 | 46.2937417 | kiosk |
| pharmacy-VIS | 10 | 9 | 2016 | 03.16.2021 | 10:00:00 | 03.29.2021 | 1 | VIS | 7.8814024 | 46.2924011 | pharmacy |
| bakery-FRI | 10 | 9 | 2016 | 03.12.21 | 09:00:00 | 03.29.2021 | 1 | FRI | 8.02164707 | 47.5069565 | bakery |
| butcher-FRI | 10 | 9 | 2016 | 03.12.21 | 09:00:00 | 03.30.2021 | 2 | FRI | 8.02079112 | 47.5075814 | butcher |

| | | | | | | | | | | | |
|---|---|---|---|---|---|---|---|---|---|---|---|
| florist-FRI | 10 | 9 | 2016 | 03.12.21 | 09:00:00 | 03.29.2021 | 1 | FRI | 8.02188618 | 47.5069733 | florist |
| kiosk-FRI | 10 | 9 | 2016 | 03.12.21 | 09:00:00 | 03.30.2021 | 2 | FRI | 8.01281878 | 47.5068387 | kiosk |
| pharmacy-FRI | 10 | 9 | 2016 | 03.12.21 | 09:00:00 | 04.01.21 | 3 | FRI | 8.01891388 | 47.5082833 | pharmacy |
| bakery-BIA | 10 | 9 | 2016 | 03.15.2021 | 10:00:00 | 03.30.2021 | 2 | BIA | 8.97059518 | 46.359882 | bakery |
| butcher-BIA | 10 | 9 | 2016 | 03.15.2021 | 10:00:00 | 03.29.2021 | 1 | BIA | 8.97256158 | 46.3535842 | butcher |
| florist-BIA | 10 | 9 | 2016 | 03.15.2021 | 10:00:00 | 03.30.2021 | 2 | BIA | 8.97168021 | 46.3578069 | florist |
| kiosk-BIA | 10 | 9 | 2016 | 03.15.2021 | 10:00:00 | 03.29.2021 | 1 | BIA | 8.97440663 | 46.3512829 | kiosk |
| pharmacy-BIA | 10 | 9 | 2016 | 03.15.2021 | 10:00:00 | 04.01.21 | 3 | BIA | 8.96984933 | 46.360162 | pharmacy |
| bakery-KLO | 10 | 9 | 2016 | 03.24.2021 | 10:00:00 | 04.01.21 | 3 | KLO | 9.88355785 | 46.8670851 | bakery |
| butcher-KLO | 10 | 9 | 2016 | 03.24.2021 | 10:00:00 | 03.30.2021 | 2 | KLO | 9.88288202 | 46.8701145 | butcher |
| florist-KLO | 10 | 9 | 2016 | 03.24.2021 | 10:00:00 | 03.29.2021 | 1 | KLO | 9.88302037 | 46.8670966 | florist |
| kiosk-KLO | 10 | 9 | 2016 | 03.24.2021 | 10:00:00 | 03.30.2021 | 2 | KLO | 9.88156669 | 46.8686036 | kiosk |
| pharmacy-KLO | 10 | 9 | 2016 | 03.24.2021 | 10:00:00 | 03.29.2021 | 1 | KLO | 9.8820039 | 46.8678293 | pharmacy |
| bakery-LAN | 10 | 9 | 2016 | 03.13.2021 | 11:00:00 | 03.29.2021 | 1 | LAN | 8.54303664 | 47.2873209 | bakery |
| butcher-LAN | 10 | 9 | 2016 | 03.13.2021 | 11:00:00 | 03.30.2021 | 2 | LAN | 8.53988255 | 47.2888713 | butcher |
| florist-LAN | 10 | 9 | 2016 | 03.13.2021 | 10:00:00 | 03.30.2021 | 2 | LAN | 8.53822798 | 47.2881315 | florist |
| kiosk-LAN | 10 | 9 | 2016 | 03.13.2021 | 11:00:00 | 03.29.2021 | 1 | LAN | 8.54375951 | 47.2871161 | kiosk |
| pharmacy-LAN | 10 | 9 | 2016 | 03.13.2021 | 10:00:00 | 04.01.21 | 3 | LAN | 8.53897478 | 47.2890869 | pharmacy |
| bakery-STE | 10 | 9 | 2016 | 03.23.2021 | 09:00:00 | 03.30.2021 | 2 | STE | 8.8585099 | 47.6594565 | bakery |
| butcher-STE | 10 | 9 | 2016 | 03.23.2021 | 09:00:00 | 04.01.21 | 3 | STE | 8.85894447 | 47.659775 | butcher |
| florist-STE | 10 | 9 | 2016 | 03.23.2021 | 09:00:00 | 03.29.2021 | 1 | STE | 8.85433058 | 47.6590038 | florist |
| kiosk-STE | 10 | 9 | 2016 | 03.23.2021 | 09:00:00 | 03.30.2021 | 2 | STE | 8.85515711 | 47.6560524 | kiosk |
| pharmacy-STE | 10 | 9 | 2016 | 03.23.2021 | 09:00:00 | 03.29.2021 | 1 | STE | 8.85932587 | 47.6595994 | pharmacy |
| bakery-ROR | 10 | 9 | 2017 | 03.18.2021 | 10:00:00 | 04.01.21 | 3 | ROR | 9.49191423 | 47.4776962 | bakery |
| butcher-ROR | 10 | 9 | 2016 | 03.18.2021 | 09:00:00 | 03.29.2021 | 1 | ROR | 9.49141016 | 47.4766705 | butcher |
| florist-ROR | 10 | 9 | 2016 | 03.18.2021 | 10:00:00 | 03.29.2021 | 1 | ROR | 9.49193047 | 47.4784607 | florist |
| kiosk-ROR | 10 | 9 | 2016 | 03.18.2021 | 10:00:00 | 03.30.2021 | 2 | ROR | 9.49274292 | 47.4785363 | kiosk |
| pharmacy-ROR | 10 | 9 | 2016 | 03.18.2021 | 10:00:00 | 03.30.2021 | 2 | ROR | 9.49109824 | 47.4778727 | pharmacy |

**Supplementary Table 3. Number, morphology and putative identity of colonies in plated samples.** The table lists the number, morphology and identity (based on identification with chromatogenic agar) of the colonies detected in plated samples (50µL aliquot on Chromatic$^{TM}$ MH agar plate, 24h at 37°C).

| Sample ID | Number of colonies | Colony morphology | Identification |
|---|---|---|---|
| bakery-ORB | 0 | n.a. | n.a. |
| butcher-ORB | 10 | green/blue, small, round, cluster | *Klebsiella, Enterobacter, Serratia* |
| florist-ORB | 2 | green/turquoise, small, round | *Enterococcus faecalis* |
| kiosk-ORB | 0 | n.a. | n.a. |
| pharmacy-ORB | 0 | n.a. | n.a. |
| bakery-POR | 0 | n.a. | n.a. |
| butcher-POR | 1 | green/blue, small, round | *Klebsiella, Enterobacter, Serratia* |
| butcher-POR | 1 | white-ish, small, round | *Staphylococcus aureus* |
| florist-POR | 0 | n.a. | n.a. |
| kiosk-POR | 0 | n.a. | n.a. |
| pharmacy-POR | 2 | green/blue, small, round | *Klebsiella, Enterobacter, Serratia* |
| pharmacy-POR | 1 | green/turquoise, small, round | *Enterococcus faecalis* |
| bakery-BRI | 2 | green/blue, small, round | *Klebsiella, Enterobacter, Serratia* |
| bakery-BRI | 1 | white-ish, small, round | *Staphylococcus aureus* |
| butcher-BRI | 5 | white-ish, small, round | *Staphylococcus aureus* |
| florist-BRI | 0 | n.a. | n.a. |
| kiosk-BRI | 0 | n.a. | n.a. |

| | | | |
|---|---|---|---|
| pharmacy-BRI | 0 | n.a. | n.a. |
| bakery-VIS | 1 | blue centre/white border, felt-ish, big (d >5mm), yeast-like | unknown |
| butcher-VIS | 14 | green/blue, small, round | *Klebsiella, Enterobacter, Serratia* |
| butcher-VIS | 2 | white-ish, small, round | *Staphylococcus aureus* |
| butcher-VIS | 59 | dark pink/red/lila, small, round | *Escherichia coli* |
| florist-VIS | 1 | dark pink/red/lila, very small, round | *Escherichia coli* |
| florist-VIS | 1 | white, translucent, big (d 4mm), yeast-like | unknown |
| kiosk-VIS | 0 | n.a. | n.a. |
| pharmacy-VIS | 0 | n.a. | n.a. |
| bakery-FRI | 1 | white-ish or light pink/red/lila, small, round | *Staphylococcus aureus* |
| butcher-FRI | 1 | dark pink/red/lila, small, round | E.coli |
| butcher-FRI | 1 | white-ish, small, round | *Staphylococcus aureus* |
| florist-FRI | 0 | n.a. | n.a. |
| kiosk-FRI | 2 | white-ish/translucent, very small, round | *Staphylococcus aureus* |
| pharmacy-FRI | 0 | n.a. | n.a. |
| bakery-BIA | 1 | yellow/white, small, 3D/growing upwards, looks like somehting you would get from your nose | unknown |
| butcher-BIA | 3 | dark pink/red/lila, small, round | *Escherichia coli* |
| butcher-BIA | 4 | white-ish, big (d 3mm), round | *Staphylococcus aureus* |
| butcher-BIA | 2 | white-ish, small, round | *Staphylococcus aureus* |
| florist-BIA | 0 | n.a. | n.a. |
| kiosk-BIA | 1 | blue, translucent, big (d 1cm), irregular shape | unknown |
| pharmacy-BIA | 0 | n.a. | n.a. |
| bakery-KLO | 0 | n.a. | n.a. |
| butcher-KLO | 13 | light pink/red/lila, small, round | *Escherichia coli* |
| butcher-KLO | 1 | green/blue, small, round | *Klebsiella, Enterobacter, Serratia* |

| | | | |
|---|---|---|---|
| butcher-KLO | 1 | green/blue, big (d >5mm), dark at centre, getting lighter going outwards, astral/yeast-like | unknown |
| florist-KLO | 0 | n.a. | n.a. |
| kiosk-KLO | 1 | green/blue, small, round | *Klebsiella, Enterobacter, Serratia* |
| kiosk-KLO | 1 | white-ish, big (d 3mm), round | *Staphylococcus aureus* |
| pharmacy-KLO | 0 | n.a. | n.a. |
| bakery-LAN | 1 | white, felt-ish, big (d 5mm), with clear bubble-like centre | unknown |
| butcher-LAN | 0 | n.a. | n.a. |
| florist-LAN | 1 | white-ish, small, round | *Staphylococcus aureus* |
| kiosk-LAN | 1 | dark pink/red/lila, small, round | *Escherichia coli* |
| pharmacy-LAN | 0 | n.a. | n.a. |
| bakery-STE | 0 | n.a. | n.a. |
| butcher-STE | 0 | n.a. | n.a. |
| florist-STE | 9 | white-ish, small, round | *Staphylococcus aureus* |
| florist-STE | 1 | light-blue, translucent, big (d 3mm), round | unknown |
| florist-STE | 2 | green/blue, small, round | *Klebsiella, Enterobacter, Serratia* |
| florist-STE | 1 | dark pink/red/lila or brown, small, round | *Escherichia coli* |
| kiosk-STE | 1 | white-ish, small, round | *Staphylococcus aureus* |
| pharmacy-STE | 0 | n.a. | n.a. |
| bakery-ROR | 0 | n.a. | n.a. |
| butcher-ROR | 1 | green/blue, small, round | *Klebsiella, Enterobacter, Serratia* |
| butcher-ROR | 1 | dark pink/red/lila, small, round | *Escherichia coli* |
| florist-ROR | 1 | yellow/green, small, round | *Pseudomonas* |
| florist-ROR | 1 | dark pink/red/lila, big (d 3mm), round | *Escherichia coli* |
| kiosk-ROR | 0 | n.a. | n.a. |

| | | | |
|---|---|---|---|
| pharmacy-ROR | 0 | n.a. | n.a. |

**Supplementary Table 4. Sequences of primers used during library preparation.** These primers were used to amplify the V3-V4 regions of the 16S rRNA gene during the first course of limited cycle PCR.

| **forward primers** | |
|---|---|
| 340F_nex0 | TCGTCGGCAGCGTCAGATGTGTATAAGAGACAGGCCTACGGGNGGCWGCAG |
| 340F_nex1 | TCGTCGGCAGCGTCAGATGTGTATAAGAGACAGNGCCTACGGGNGGCWGCAG |
| 340F_nex2 | TCGTCGGCAGCGTCAGATGTGTATAAGAGACAGNNGCCTACGGGNGGCWGCAG |
| 340F_nex3 | TCGTCGGCAGCGTCAGATGTGTATAAGAGACAGNNNGCCTACGGGNGGCWGCAG |
| **reverse primers** | |
| 805R_nex0 | GTCTCGTGGGCTCGGAGATGTGTATAAGAGACAGTGACTACHVGGGTATCTAATCC |
| 805R_nex1 | GTCTCGTGGGCTCGGAGATGTGTATAAGAGACAGNTGACTACHVGGGTATCTAATCC |
| 805R_nex2 | GTCTCGTGGGCTCGGAGATGTGTATAAGAGACAGNNTGACTACHVGGGTATCTAATCC |
| 805R_nex3 | GTCTCGTGGGCTCGGAGATGTGTATAAGAGACAGNNNTGACTACHVGGGTATCTAATCC |

**Supplementary Table 5. Reads per sample before and after filtering.** The table lists the read sums for each sample before and after each filtering step. Filtering step 1 (ReadSum_01 → ReadSum_02): removal of samples with read sum <1000 (n = 8, florist-KLO, butcher-POR, florist-POR, bakery-STE, florist-STE, bakery-BIA, bakery-VIS, florist-LAN). Filtering step 2 (ReadSum_02 → ReadSum_03): removal of ZOTUs present at <0.1% of total sequencing depth (results in reduction in read sum for each sample; total sequencing depth = 14’398’160). The location abbreviations are: BIA = Biasca, BRI = Brienz BE, FRI = Frick, KLO = Klosters-Serneus, LAN = Langnau am Albis, ORB = Orbe, POR = Porrentruy, ROR = Rorschach, STE = Stein am Rhein, VIS = Visp.

| Sample ID | ReadSum_01 | ReadSum_02 | ReadSum_03 | Location | Shop type |
|---|---|---|---|---|---|
| bakery-FRI | 114200 | 114200 | 53760 | FRI | bakery |
| butcher-FRI | 162322 | 162322 | 93660 | FRI | butcher |
| florist-FRI | 202155 | 202155 | 42515 | FRI | florist |
| kiosk-FRI | 175437 | 175437 | 85868 | FRI | kiosk |
| pharmacy-FRI | 307244 | 307244 | 61859 | FRI | pharmacy |
| bakery-BRI | 5328 | 5328 | 3392 | BRI | bakery |
| butcher-BRI | 2060 | 2060 | 138 | BRI | butcher |
| florist-BRI | 6752341 | 6752341 | 6167513 | BRI | florist |
| kiosk-BRI | 208071 | 208071 | 198031 | BRI | kiosk |
| pharmacy-BRI | 9225 | 9225 | 6040 | BRI | pharmacy |
| bakery-KLO | 202364 | 202364 | 98592 | KLO | bakery |
| butcher-KLO | 235452 | 235452 | 179502 | KLO | butcher |
| florist-KLO | 190 | NA | NA | KLO | florist |
| kiosk-KLO | 237475 | 237475 | 118867 | KLO | kiosk |
| pharmacy-KLO | 191824 | 191824 | 97575 | KLO | pharmacy |
| bakery-POR | 308617 | 308617 | 178351 | POR | bakery |
| butcher-POR | 4 | NA | NA | POR | butcher |
| florist-POR | 219 | NA | NA | POR | florist |
| kiosk-POR | 320526 | 320526 | 257093 | POR | kiosk |
| pharmacy-POR | 345879 | 345879 | 147741 | POR | pharmacy |
| bakery-ROR | 299698 | 299698 | 179789 | ROR | bakery |
| butcher-ROR | 95170 | 95170 | 50599 | ROR | butcher |
| florist-ROR | 181294 | 181294 | 25490 | ROR | florist |
| kiosk-ROR | 305964 | 305964 | 153294 | ROR | kiosk |
| pharmacy-ROR | 114889 | 114889 | 39322 | ROR | pharmacy |
| bakery-STE | 166 | NA | NA | STE | bakery |
| butcher-STE | 324269 | 324269 | 149241 | STE | butcher |
| florist-STE | 290 | NA | NA | STE | florist |
| kiosk-STE | 94564 | 94564 | 36712 | STE | kiosk |
| pharmacy-STE | 374098 | 374098 | 171063 | STE | pharmacy |
| bakery-BIA | 23 | NA | NA | BIA | bakery |

| | | | | | |
|---|---|---|---|---|---|
| butcher-BIA | 281304 | 281304 | 213667 | BIA | butcher |
| florist-BIA | 68992 | 68992 | 68351 | BIA | florist |
| kiosk-BIA | 375140 | 375140 | 231005 | BIA | kiosk |
| pharmacy-BIA | 1331 | 1331 | 758 | BIA | pharmacy |
| bakery-ORB | 102777 | 102777 | 90516 | ORB | bakery |
| butcher-ORB | 154164 | 154164 | 63229 | ORB | butcher |
| florist-ORB | 65900 | 65900 | 15267 | ORB | florist |
| kiosk-ORB | 212692 | 212692 | 79226 | ORB | kiosk |
| pharmacy-ORB | 2775 | 2775 | 838 | ORB | pharmacy |
| bakery-VIS | 722 | NA | NA | VIS | bakery |
| butcher-VIS | 302458 | 302458 | 238540 | VIS | butcher |
| florist-VIS | 343588 | 343588 | 54478 | VIS | florist |
| kiosk-VIS | 167685 | 167685 | 94603 | VIS | kiosk |
| pharmacy-VIS | 223900 | 223900 | 118012 | VIS | pharmacy |
| bakery-LAN | 88027 | 88027 | 72522 | LAN | bakery |
| butcher-LAN | 258637 | 258637 | 129820 | LAN | butcher |
| florist-LAN | 18 | NA | NA | LAN | florist |
| kiosk-LAN | 2586 | 2586 | 1093 | LAN | kiosk |
| pharmacy-LAN | 175738 | 175738 | 68254 | LAN | pharmacy |

**Supplementary Table 6. Relative abundance of the top 12 families across samples.** The table lists the relative abundance (in %) of the 12 relatively most abundant families for all samples. The location abbreviations are: BIA = Biasca, BRI = Brienz BE, FRI = Frick, KLO = Klosters-Serneus, LAN = Langnau am Albis, ORB = Orbe, POR = Porrentruy, ROR = Rorschach, STE = Stein am Rhein, VIS = Visp. Latitude and longitude are given in WGS format.

Due to the size of the file this table is available separately (*suppTable_06.csv*).

**Supplementary Table 7. Relative abundance of the top 12 genera across samples.** The table lists the relative abundance (in %) of the 12 relatively most abundant genera for all samples. The location abbreviations are: BIA = Biasca, BRI = Brienz BE, FRI = Frick, KLO = Klosters-Serneus, LAN = Langnau am Albis, ORB = Orbe, POR = Porrentruy, ROR = Rorschach, STE = Stein am Rhein, VIS = Visp. Latitude and longitude are given in WGS format.

Due to the size of the file this table is available separately (*suppTable_07.csv*).

**Supplementary Table 8. Relative abundance of shared core ZOTUs.** The table lists the relative abundance (in %) of the seven dominant core ZOTUs (Dawson *et al.*, 2017; Cruaud *et al.*, 2020) shared between all shop types. Location abbreviations are: BIA = Biasca, BRI = Brienz BE, FRI = Frick, KLO = Klosters-Serneus, LAN = Langnau am Albis, ORB = Orbe, POR = Porrentruy, ROR = Rorschach, STE = Stein am Rhein, VIS = Visp.

Due to the size of the file this table is available separately (*suppTable_08.csv*).

**Supplementary Table 9. NMDS scores for each of the 163 ZOTUs.** The table lists the scores for each dimension (NMDS1, NMDS2) in the ordination visualisation (Fig. 3). ZOTU annotation with format_to_besthit() function in microbiomeutilities package (version 1.00.16; (Lahti *et al.*, 2017)).

| ZOTU | MDS1 | MDS2 | ZOTU | MDS1 | MDS2 |
|---|---|---|---|---|---|
| *ZOTU2:Escherichia-Shigella* | 0.05730727 | -0.0327834 | *ZOTU142:Bradyrhizobiaceae* | 0.01442155 | -0.0724633 |
| *ZOTU41:Streptococcus* | 0.05328139 | -0.0106425 | *ZOTU172:Corynebacteriaceae* | 0.01340052 | -0.0172705 |
| *ZOTU56:Veillonella* | 0.05276656 | 0.00339363 | *ZOTU79:Nocardioides* | 0.01304672 | 0.01741828 |
| *ZOTU140:Rothia* | 0.05274238 | -0.0256805 | *ZOTU11:Saccharibacteria* | 0.0125994 | -0.0211877 |
| *ZOTU78:Rothia* | 0.05192814 | -0.0129222 | *ZOTU111:Desemzia* | 0.01134844 | -0.0078049 |
| *ZOTU1:Pseudomonas* | 0.05148414 | -0.0267068 | *ZOTU99:Corynebacteriaceae* | 0.01024344 | 0.08483568 |
| *ZOTU87:Streptococcus* | 0.05132028 | -0.0123853 | *ZOTU121:Lactococcus* | 0.01000869 | 0.13337945 |
| *ZOTU3:Staphylococcus* | 0.05095895 | -0.0171798 | *ZOTU31:Thermus* | 0.00904807 | -0.0307946 |
| *ZOTU38:Propionibacterium* | 0.05081344 | -0.0137691 | *ZOTU33:Flavobacterium* | 0.00849264 | -0.0176427 |
| *ZOTU5:Streptococcus* | 0.0505481 | -0.0118718 | *ZOTU169:Clostridiisalibacter* | 0.00797463 | 0.02272327 |
| *ZOTU21:Rothia* | 0.05037946 | -0.0130542 | *ZOTU55:Photobacterium* | 0.00721648 | 0.09179588 |
| *ZOTU88:Corynebacteriaceae* | 0.05033086 | -0.0269886 | *ZOTU46:Lactobacillales* | 0.00709748 | -0.0191396 |
| *ZOTU6:Corynebacterium* | 0.05028706 | -0.0113914 | *ZOTU92:Lactobacillus* | 0.00586005 | 0.08017116 |
| *ZOTU64:Streptococcus* | 0.04959371 | 0.01789076 | *ZOTU107:Sphingomonas* | 0.00531963 | -0.0277524 |
| *ZOTU161:Lawsonella* | 0.04943638 | -0.0271418 | *ZOTU163:Moraxella_sp._feline _oral_taxon_089* | 0.00510093 | -0.0877364 |
| *ZOTU139:Granulicatella* | 0.04937002 | -0.0219911 | *ZOTU166:Chloroflexi* | 0.0033126 | -0.0779263 |
| *ZOTU89:Brochothrix* | 0.04828398 | 0.03054945 | *ZOTU162:Leuconostoc* | 0.0024969 | 0.10316701 |
| *ZOTU20:Gemella* | 0.04818861 | -0.0319006 | *ZOTU104:Corynebacterium* | 0.00047779 | -0.0307464 |
| *ZOTU174:Sphingomonas* | 0.04789879 | -0.0110251 | *ZOTU153:Photobacterium* | -0.0006998 | 0.13064208 |
| *ZOTU13:Staphylococcus* | 0.04664732 | 0.01109724 | *ZOTU160:Streptococcus* | -0.002533 | 0.00634604 |
| *ZOTU117:Rothia* | 0.04656358 | -0.0133407 | *ZOTU47:Brevibacterium* | -0.0028092 | 0.05083274 |
| *ZOTU4:Streptococcus* | 0.04601654 | -0.0025 | *ZOTU68:Enterobacteriaceae* | -0.0031899 | -0.03016 |
| *ZOTU44:Novosphingobium* | 0.04571736 | -0.013307 | *ZOTU168:Aerococcus* | -0.0036755 | 0.03272319 |
| *ZOTU14:Pseudomonas* | 0.04559882 | -0.0201748 | *ZOTU50:Photobacterium* | -0.0116215 | 0.20624871 |
| *ZOTU196:Corynebacterium* | 0.04540179 | -0.0159044 | *ZOTU74:Brevibacterium* | -0.0151122 | 0.10541356 |
| *ZOTU120:Haemophilus* | 0.04434457 | -0.002158 | *ZOTU30:Brevibacterium* | -0.0179297 | 0.08180127 |
| *ZOTU39:Enhydrobacter* | 0.04415742 | -0.0152317 | *ZOTU17:Pseudarthrobacter* | -0.0205082 | -0.007437 |
| *ZOTU49:Staphylococcus* | 0.04411697 | 0.02237678 | *ZOTU98:Brevibacterium* | -0.0225247 | 0.10207496 |
| *ZOTU201:Corynebacterium* | 0.04313749 | 0.00594622 | *ZOTU127:Lawsonella* | -0.030283 | -0.0347926 |
| *ZOTU108:Anaerococcus* | 0.04257978 | 0.0075745 | *ZOTU138:Planifilum* | -0.0321269 | -0.0760371 |
| *ZOTU16:Staphylococcus* | 0.0424552 | -0.0101929 | *ZOTU134:Duganella* | -0.040449 | 0.06389764 |
| *ZOTU143:Abiotrophia* | 0.04154395 | -0.0109499 | *ZOTU23:Acidimicrobiales* | -0.0422583 | -0.0423177 |
| *ZOTU179:Psychrobacter* | 0.04089985 | -0.010511 | *ZOTU148:Lysobacter* | -0.0442132 | -0.0172773 |
| *ZOTU73:Kocuria* | 0.0406755 | 0.00154943 | *ZOTU18:Acetobacteraceae* | -0.0553217 | -0.0386851 |

| | | | | | |
|---|---|---|---|---|---|
| *ZOTU129:Paracoccus* | 0.04019151 | 0.00169117 | *ZOTU15:Rhizobiales* | -0.0610517 | -0.0337397 |
| *ZOTU28:Micrococcus* | 0.04012933 | -0.0010915 | *ZOTU22:Parcubacteria* | -0.0648825 | -0.034022 |
| *ZOTU137:Truepera* | 0.03992759 | -0.0462813 | *ZOTU173:Cellulomonadaceae* | -0.0698148 | -0.0404219 |
| *ZOTU109:Gemella* | 0.03986634 | -0.0276838 | *ZOTU40:Anaerococcus* | -0.0733416 | -0.0519969 |
| *ZOTU36:Chryseobacterium* | 0.03854048 | -0.0029423 | *ZOTU76:Gaiella* | -0.0734224 | -0.0279952 |
| *ZOTU91:Micrococcus* | 0.03842443 | 0.0044853 | *ZOTU54:Corynebacteriaceae* | -0.0762838 | 0.15066023 |
| *ZOTU29:Porphyromonas* | 0.0367369 | -0.0216445 | *ZOTU52:Sphingobacterium* | -0.0968468 | 0.08088874 |
| *ZOTU133:Rhodococcus* | 0.03664773 | 0.0133623 | *ZOTU167:Mycobacterium* | -0.1082307 | 0.00501021 |
| *ZOTU197:Veillonella* | 0.03589104 | 0.0294119 | *ZOTU61:Photobacterium* | -0.1097046 | 0.39919256 |
| *ZOTU178:Acinetobacter* | 0.03515831 | 0.01292604 | *ZOTU170:Kocuria* | -0.1159726 | 0.12922351 |
| *ZOTU125:Lactococcus* | 0.03512806 | -0.0099691 | *ZOTU152:Isosphaera* | -0.1162278 | -0.0735077 |
| *ZOTU34:Staphylococcus* | 0.03496982 | 0.01011052 | *ZOTU115:Microbacteriaceae* | -0.1163409 | 0.05872633 |
| *ZOTU37:Staphylococcus* | 0.03492365 | -0.0137789 | *ZOTU128:Hymenobacter* | -0.1316927 | -0.1088893 |
| *ZOTU102:Finegoldia* | 0.03362732 | -0.0045215 | *ZOTU124:Actinomycetospora* | -0.1319739 | -0.0187272 |
| *ZOTU103:Carnobacterium* | 0.03352972 | 0.0368222 | *ZOTU100:Photobacterium* | -0.1358745 | 0.34840205 |
| *ZOTU118:Neisseriaceae* | 0.03333235 | -0.0375732 | *ZOTU165:Halomonas* | -0.1897639 | 0.2433281 |
| *ZOTU187:Craurococcus* | 0.03328822 | -0.0640929 | *ZOTU84:Flavobacterium* | -0.2049448 | -0.0463485 |
| *ZOTU146:Staphylococcus* | 0.03184716 | -0.0263628 | *ZOTU176:Brachybacterium* | -0.2060346 | 0.3107196 |
| *ZOTU32:Micrococcaceae* | 0.03132497 | -0.0046581 | *ZOTU171:Methylobacterium* | -0.2390657 | 0.01347099 |
| *ZOTU70:Leuconostoc* | 0.03102254 | 0.05085632 | *ZOTU94:Lachnospiraceae* | -0.2484112 | 0.01217332 |
| *ZOTU80:Acinetobacter* | 0.03087387 | 0.00694619 | *ZOTU112:Hymenobacter* | -0.3967924 | -0.0092792 |
| *ZOTU101:Peptoniphilus* | 0.03010505 | -0.0629798 | *ZOTU63:OM1_clade* | -0.4748172 | -0.1453787 |
| *ZOTU77:Acidovorax* | 0.02962544 | -0.031069 | *ZOTU72:Subdoligranulum* | -0.5410566 | -0.0019624 |
| *ZOTU131:Acinetobacter* | 0.02935586 | 0.0410873 | *ZOTU122:Gaiellaceae* | -0.576009 | -0.1866043 |
| *ZOTU86:Streptococcus* | 0.02928791 | -0.0180964 | *ZOTU110:Singulisphaera* | -0.6027149 | -0.0591442 |
| *ZOTU45:Corynebacterium* | 0.0291601 | -0.0356454 | *ZOTU51:Variibacter* | -0.6044048 | 0.00423743 |
| *ZOTU82:Corynebacterium_1* | 0.02898465 | 0.04453783 | *ZOTU147:Pseudomonas* | -0.6050551 | -0.1061542 |
| *ZOTU19:Corynebacteriaceae* | 0.02894523 | -0.0241684 | *ZOTU48:Stenotrophomonas* | -0.6072141 | -0.0522472 |
| *ZOTU8:Corynebacterium* | 0.02771125 | 0.00296455 | *ZOTU208:Bacilli* | -0.6102026 | -0.2618254 |
| *ZOTU26:Bacillus* | 0.02751467 | -0.0350773 | *ZOTU43:Alphaproteobacteria* | -0.617965 | -0.0669666 |
| *ZOTU66:Microbacteriaceae* | 0.02702594 | -0.0200106 | *ZOTU149:Porphyromonas* | -0.6389732 | -0.1987567 |
| *ZOTU69:Lactobacillus* | 0.02634711 | 0.07936559 | *ZOTU25:Acidiphilium* | -0.6904041 | -0.0956538 |
| *ZOTU114:Staphylococcus* | 0.02633792 | 0.0349737 | *ZOTU157:Subgroup* | -0.6976051 | 0.08155747 |
| *ZOTU119:Rothia* | 0.02466601 | 0.01108826 | *ZOTU105:Bryobacter* | -0.7101991 | -0.1281004 |
| *ZOTU154:Methylobacterium* | 0.02415951 | -0.0654888 | *ZOTU62:Tepidisphaeraceae* | -0.7166585 | -0.0277752 |
| *ZOTU158:Brevibacterium* | 0.0235347 | 0.00785889 | *ZOTU71:Acidimicrobiaceae* | -0.7317721 | -0.084418 |
| *ZOTU126:Oxalobacteraceae* | 0.02153712 | -0.0223155 | *ZOTU136:Ktedonobacteraceae* | -0.7926883 | -0.0885217 |
| *ZOTU58:Staphylococcus* | 0.02100687 | -0.0385819 | *ZOTU141:Chitinophagaceae* | -0.7999002 | -0.0702119 |
| *ZOTU96:Kocuria* | 0.01949318 | 0.04142059 | *ZOTU130:Saccharibacteria* | -0.8013313 | -0.0645734 |

| | | | | | |
|---|---|---|---|---|---|
| *ZOTU42:Intrasporangiaceae* | 0.01923794 | 0.00358067 | *ZOTU195:Hymenobacter* | -0.8096099 | -0.079094 |
| *ZOTU192:Pseudomonas* | 0.01880303 | 0.00183501 | *ZOTU116:Hymenobacter* | -0.8183641 | -0.0809633 |
| *ZOTU113:Dolosigranulum* | 0.01840913 | -0.0126963 | *ZOTU159:Ktedonobacteraceae* | -0.83517 | -0.0985353 |
| *ZOTU210:Actinomyces* | 0.01828953 | -0.0091106 | *ZOTU83:Hymenobacter* | -0.8380494 | -0.0762446 |
| *ZOTU151:Massilia* | 0.01751111 | -0.0482222 | *ZOTU90:Desemzia* | -0.8410246 | -0.0767683 |
| *ZOTU24:Kocuria* | 0.01721711 | -0.0040575 | *ZOTU95:Saccharibacteria* | -0.8528812 | -0.0802721 |
| *ZOTU12:Turicella* | 0.01719001 | -0.0002162 | | | |
| *ZOTU97:Flavobacterium* | 0.01683197 | -0.016063 | | | |
| *ZOTU60:Comamonadaceae* | 0.01464276 | -0.0490035 | | | |
| *ZOTU59:Staphylococcus* | 0.01459722 | -0.0427289 | | | |
| *ZOTU145:Intrasporangiaceae* | 0.01459563 | 0.0033087 | | | |
